# Cell division angle regulates the tissue mechanics and tunes the amount of cerebellar folding

**DOI:** 10.1101/2023.07.21.549165

**Authors:** Amber G. Cook, Taylor V. Bishop, Hannah R. Crowe, Daniel Stevens, Lauren Reine, Alexandra L Joyner, Andrew K Lawton

## Abstract

Modeling has proposed that the amount of neural tissue folding is set by the level of differential-expansion between tissue layers and that the wavelength is set by the thickness of the outer layer. Here we used inbred mouse strains with distinct amounts of cerebellar folding to investigate these predictions. We identified a critical period where the folding amount diverges between the strains. In this period, regional changes in the level of differential-expansion between the external granule layer (EGL) and underlying core correlate with the folding amount in each strain. Additionally, the thickness of the EGL is regionally adjusted during the critical period alongside corresponding changes in wavelength. While the number of SHH-expressing Purkinje cells predicts the folding amount, the proliferation rate in the EGL is the same between the strains. However, regional changes in the cell division angle within the EGL predicts both the tangential-expansion and thickness of the EGL. Cell division angle is likely a tunable mechanism whereby both the level of differential-expansion and thickness of the EGL are regionally tuned to set the amount and wavelength of folding.

## Introduction

The human cortex and cerebellum are formed into complex folded geometries that provide space for the substantial neural circuits and a gross 3-D compartmentalization of functional circuitry within each structure. Changes to the amount of folding in the brain are associated with intellectual disabilities, epilepsies and other health issues (Flotats-Bastardas et al., 2012; Leventer et al., 2010; Raymond et al., 1995). Therefore, determining how the degree of folding is set during development is critical for understanding the functional role of the 3-D partitioning of circuits in the gyri of the cortex and the lobules of the cerebellum.

Little is known about how the tissue geometries and mechanics work together to set the proper amount of folding during development. The murine cerebellum has 8-10 folds aligned in the anterior-posterior axis in the medial region (vermis). This simple arrangement allows for a precise analysis of the geometric and mechanical regulation of the degree of folding from analysis of sagittal sections. Neural tissues have different geometries and conformations than other tissues that undergo folding-like events during development (Nelson, 2016; Shyer et al., 2013; Varner and Nelson, 2014). In the developing brain cells are arraigned, not in simple epithelial layers, but in dynamic and thick laminar structures. The cerebellum is further unique. During the second phase of cerebellar development, it grows through the rapid expansion of a temporary external granule layer (EGL) that covers its entire outer surface. The granule cell progenitors (GCPs) within the EGL are proliferative, motile, and are responsible for most of the cerebellar volume growth and for directing growth primarily in the anterior-posterior axis (Joyner and Bayin, 2022; Lawton et al., 2023; Lawton et al., 2019; Legué et al., 2016, 2015; Leto et al., 2016).

Previously we demonstrated that cerebellar folding is initiated when the ratio of growth between the EGL and the underlying core diverges enough to create a differential-expansion with the EGL growing faster (Lawton et al., 2019). This differential-expansion provides the driving force of initial cerebellar folding. Further we showed how the geometry of the unfolded tissue sets the ratio of growth required to create the differential-expansion needed for folding. These tissue mechanics, along with tissue tension and the fluid-like behavior of the EGL are predicted to fold and shape the cerebellum during embryonic development (Engstrom et al., 2018). However, these ideas have not been tested.

One prediction concerning the regulation of the folding amount is that the number of folds is set by the level of differential-expansion between the outer layer, the EGL, and the underlying cerebellar core. A second is that the wavelength of folding is defined by the thickness of the EGL (Engstrom et al., 2018; Lawton et al., 2019; Tallinen et al., 2016, 2014). Excitingly, the folding amount in the adult cerebral cortex across many species correlates with the thickness of the cerebral cortex which is its corresponding outer layer during folding (Mota and Herculano-Houzel, 2015). Further, the variation of folding within individual cerebral cortices also follows this same relationship (Wang et al., 2019). This led us to ask if the amount of cerebellar folding is set during development by adjusting both the amount of differential-expansion between the EGL and core and the thickness of the EGL.

Here we used a gene-agnostic approach to assess the tissue mechanics and tissue geometries predicted to regulate the amount of folding during development. We utilized two inbred strains of mice with small, but robust differences in the final number of vermis folds. We report that the level of differential-expansion between the EGL and core during development is regionally regulated to set the degree of folding as predicted. Further we demonstrate that the thickness of the EGL also regionally varies and is consistent with the observed wavelength. Differences in the amount of the tangential-expansion of the EGL and the lobule geometry appear to work together to regionally define the level of differential-expansion between the EGL and core. We show that regional differences in the cell-division angle within the EGL predicts the tangential-expansion of the EGL and its thickness. Thus, cell division angle could be a tunable mechanism whereby both the level of differential-expansion and the thickness of the EGL are precisely tuned to set the amount of folding in the cerebellum.

## Results

### Inbred mouse strains have regionally distinct levels of folding

We used the C57Bl/6J and FVB/NJ inbred strains of mice, since they have a robust difference in the amount of cerebellar foliation (Sillitoe and Joyner, 2007). Here we considered P28 an “adult” stage as the cerebellum has reached its final size and folding amount. In both strains most cerebella have normal cytoarchitectural layers (Fig 1 A,B, Supplemental Fig 1A-C) and are both considered “wild-type.” A subset of C57Bl/6J mice, however, have heterotopia which disturbs the pattern of folding in the posterior cerebellum (Supplemental Fig 1 B) (Van Dine et al., 2015). Individuals or lobules showing malformation were excluded from this study.

**Figure 1:**
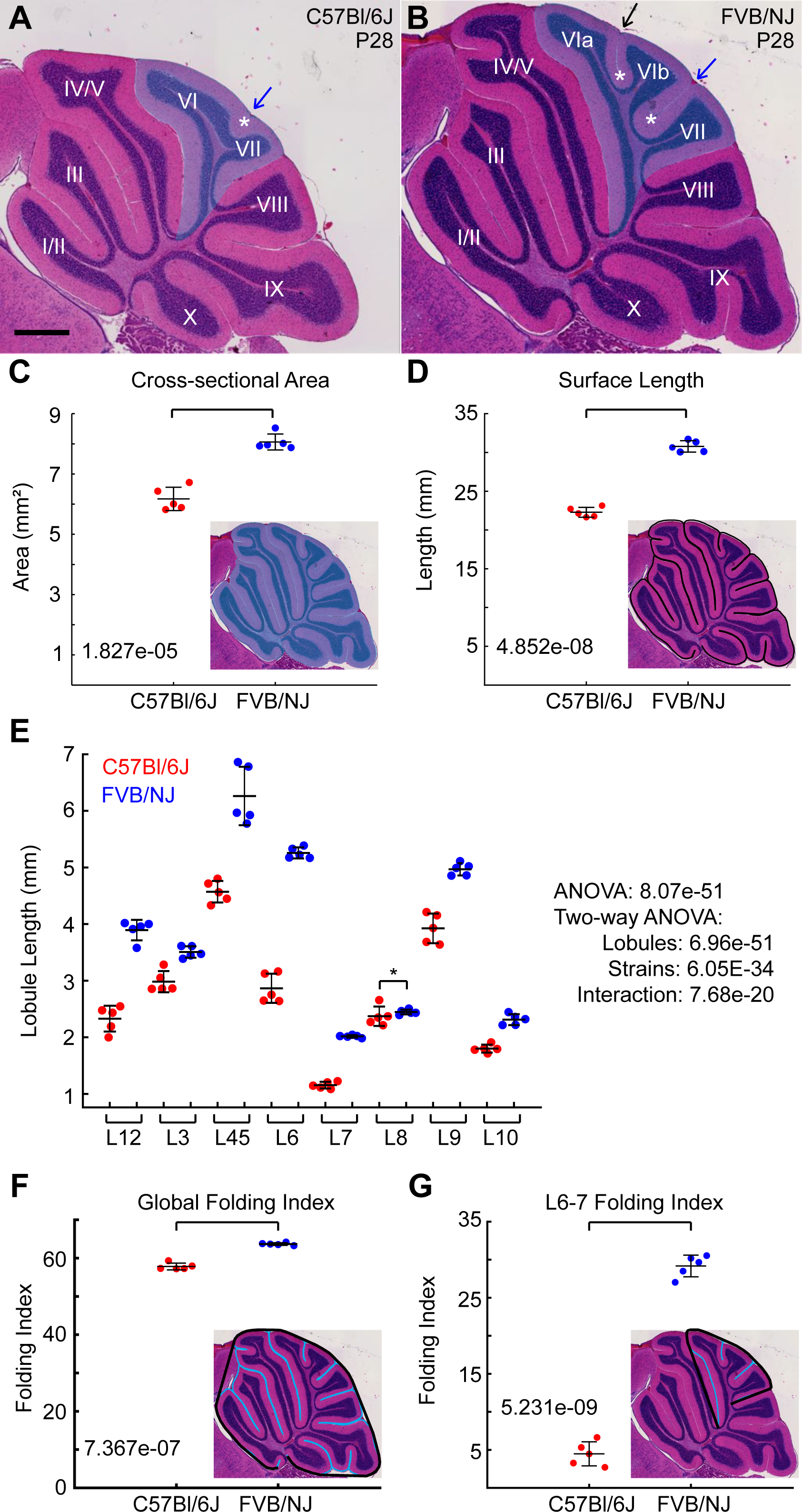
Mouse strains have regionally varied levels of cerebellar folding. **A,B**) H&E stained sagittal midline sections of P28 C57Bl/6J and FVB/NJ cerebella. Shaded region: lobules 6 and 7. Black arrow: superior-declive fissure (SDF), blue arrow: Intercrural fissure (IF) Asterisks: anchoring centers. Scale bar: 500 µM. **C**) Cross-sectional area. **D**) Surface length. **E)** All Lobule lengths are reduced in C57Bl/6J except for L8. **F**) Global folding index. **G**) Regional folding index of L6-7. **A**ll analyses: n = 5/strain (mean s.d.)

The C57Bl/6J cerebellum is significantly smaller than the FVB/NJ, in its cross-sectional area, surface length, and positive curvature (Fig 1 C, D Supplemental Fig 2A). Globally C57Bl/6J is about ∼72-75% the size of FVB/NJ in surface length and area. Dividing the cerebellum into the lobules that are fully separated in both genotypes revealed that the size decrease in C57Bl/6J was not uniform, but regionally adjusted across the cerebellum. Lobules 6 and 7 (L6-7) were strongly decreased in length whereas L8 was maintained in size compared to FVB/NJ (Fig 2E). We next measured the lobule lengths as a percentage of the total length of the cerebellum. While several lobules are proportionally scaled down in C57Bl/6J compared to FVB/NJ (L4-5, 7, 9, 10), some are increased (L3, 8) and some are decreased (L3, 6) (Supplemental Fig 2B).

**Figure 2:**
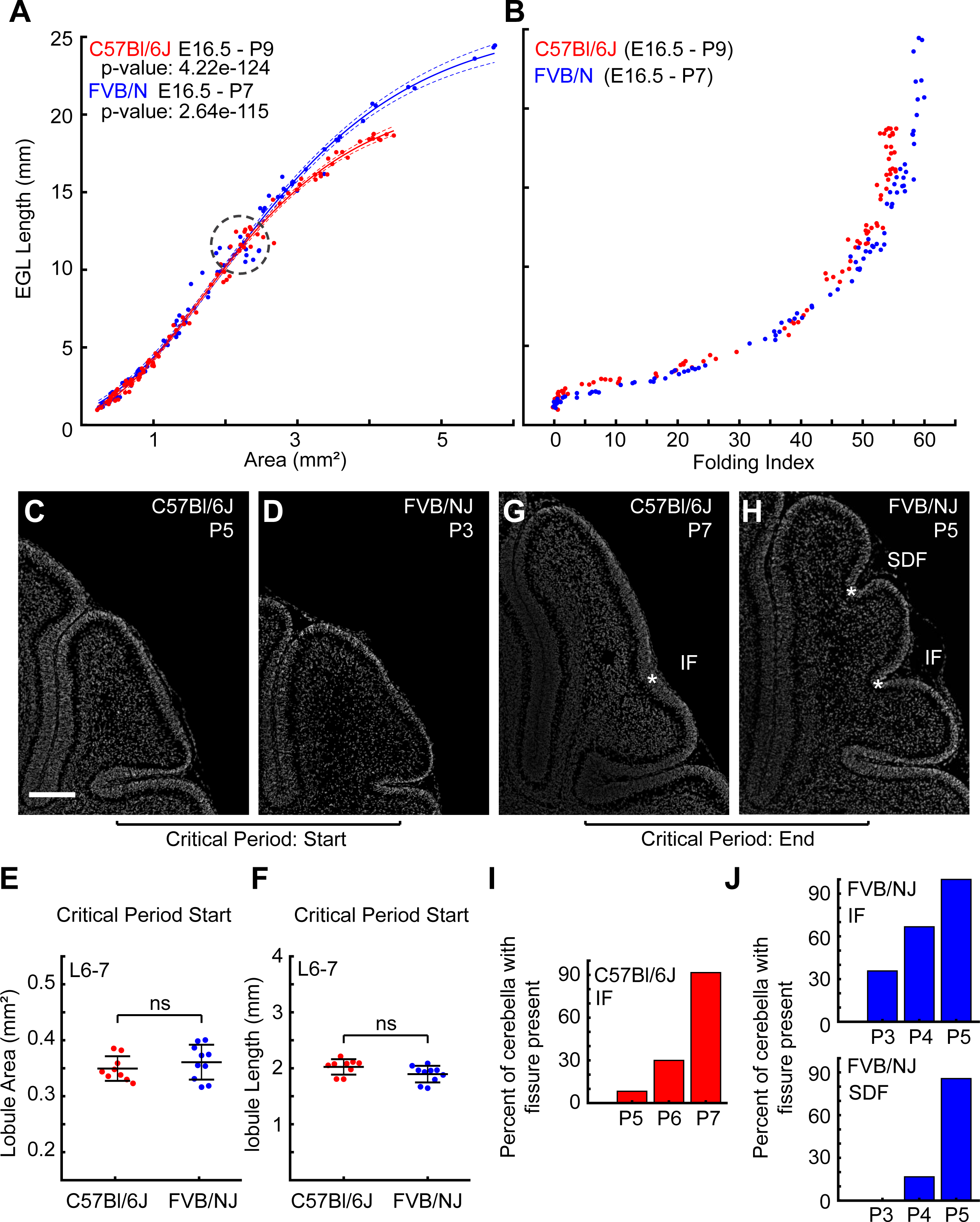
Folding differences arise during postnatal period. **A**) Growth ratios of the EGL length and cerebellar area from E16.5 through the formation of the final fissures. Lines: Gompertz fitting with confidence intervals. FVBN: n = 96 C57Bl6J: n = 94. A subset of FVB/N data below 1mm^2^ was previously published (Lawton et al., 2019). **B**) Folding index. **C,D**) L6-7 regions at the start of the critical period. Scale bar: 200µM **E,F**) L6-7 region has the same lobule area and length between strains at the start of the critical period. n = 9 C57Bl/6J; 10 FVB/NJ. **G,H**) L6-7 regions at the end of the critical period showing acquisition of the intercrural (IF) and the superior-declive (SDF) fissures. Asterisk: anchoring centers at the base of each fissure. **I-J**) Percentage of cerebella showing anchoring center acquisition. C57Bl/6J: P5: 1/12, P6: 3/10, P7: 11/12; FVB/NJ PSF P3: 5/14, P4: 8/12, P5: 14/14; FVBN/J ICF: P3: 0/14, P4: 2/12, P5: 12/14. (mean s.d.)

We measured the folding index [FI = (1-(Positive Curvature/Surface Length))*100)] to quantify the amount of folding (Lawton et al., 2019). The C57Bl/6J cerebellum is less folded than the FVB/NJ showing a ∼10% reduction (Fig 1F). However, the regional folding of the L6-7 region is reduced by around 85% (Fig 1G). In this region the C57Bl/6J lacks the superior-declive fissure (SDF) separating sub lobules 6a and 6b and the depth of the intercural fissure (IF) separating sub L6b from L7 is strongly reduced (black and blue arrows, respectively in Fig 1A,B). The folding index of the anterior (L1-5) was reduced by ∼ 8% and the posterior (L8-10) region of the cerebellum showed a ∼13% reduction in C57Bl/6J (Supplemental Fig 2C,D). The robust folding difference in the L6-7 region in these two strains provide a tractable system to test if differential-expansion and EGL thickness could be responsible for regulating folding amount during development.

### The growth ratios diverge at the onset of folding differences

Differential-expansion can emerge between tissue layers that expand at different rates (Hannezo et al., 2012; Nelson, 2016; Shyer et al., 2013; Wiggs et al., 1997). Increasing or decreasing the levels of differential-expansion is predicted to change the amount of folding (Tallinen et al., 2014). While the cerebellum has several layers and has been treated as a tri-layer developing system (Lejeune et al., 2016), the cerebellum is not separated into all the distinct cytological layers seen in the adult during the early stages of folding. The full suite of cellular layers progressively become more distinct as folding continues after birth. The first folds arise embryonically when the only distinct layers are the EGL and the underlying core. Therefore, we have treated the cerebellum as functionally behaving as a two-layer system: the outer and highly proliferative EGL and the inner core.

We measured the ratio between the rate of midsagittal EGL length increase and the rate of increase in the area from Embryonic day 16.5 (E16.5) through the stage when the final anchoring centers, the bases of the fissures, form (Fig 2A). This allowed an unbiased comparison of the growth of the strains, controlling for any global differences in the rates of development of the different strains, actual ages of the various litters due to mating time variation, and growth affects from litter size. The global growth curves are well fitted by a basic Gompertz function (Tjørve and Tjørve, 2017). This function was chosen for its minimal variables, its historic use in modeling biological growth, and how well it models the complex data. The non-linear regression analyses for the individual strains show a better fit with more uniform residuals and distinct parameters for each strain than with a combined fit (Supplemental Fig 3A-E).

We found that both C57Bl/6J and FVB/NJ had the same ratio of growth between the EGL expansion and the area expansion until they reach a size of over ∼ 2mm^2^. Correspondingly, during this early period the folding indices for each strain were overlapped as well (Fig 2B). This result indicates that the early growth ratios of the EGL and core and tissue geometries are likely creating similar amounts of differential-expansion which are in turn creating similar amounts of folding. Therefore, mechanical differences likely arise after the cerebella reach this critical point.

Interestingly, the C57Bl/6J and FVB/NJ cerebella reach the critical size at different ages, approximately P5 and P3 respectively. This finding demonstrates that FVB/NJ cerebella grow more rapidly than C57Bl/6J, to reach the same size roughly two days earlier. However, the critical factor for determining the final amount of folding is not the global speed of growth but likely the level of differential-expansion that emerges from the growth ratio between the EGL and the core area.

As the cerebella expand beyond ∼2mm^2^, the surface length of the FVB/NJ EGL continues to expand at the same rate whereas the C57Bl/6J EGL reduces its rate in relation to core area growth (Fig 2A). Once the growth ratios diverged, the amount of folding in the cerebellum also diverged (Fig. 2B). Furthermore, in both strains, when the cerebella are ∼2mm^2^, the L6-7 region is unfolded and has the same size in both EGL length and lobule area (Fig 3 C-F). However, as the global growth ratios diverged, the difference in the number of folds in the L6-7 region (one in C57Bl/6J and 2 in FVB/NJ) become apparent (Fig 2G,H). We identified a critical period of development for the L6-7 region when the SDF and IF appear and the folding amount between the strains diverges. This period coincides with an age of ∼P5-P7 for C57Bl/6J and ∼P3-P5 for FVB/NJ (Fig 2I,J).

**Figure 3.**
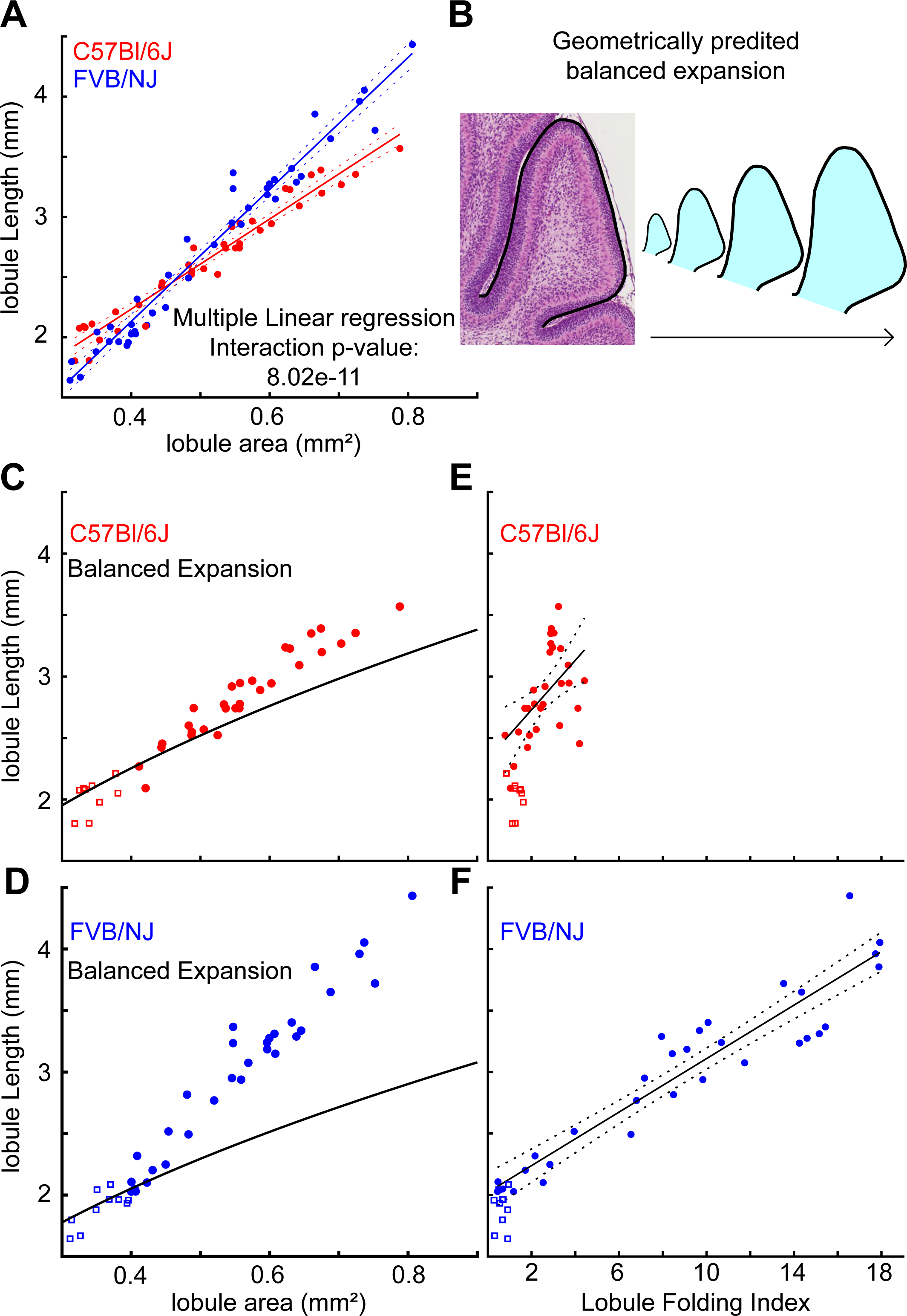
The level of differential expansion sets the folding amount. **A**) Growth ratios of L6-7 regions from the critical period. **B**) Balanced expansion curve from geometry of lobule regions at start of critical period (squares in C-F). **C**) The C57Bl/6J growth ratio remains near balanced growth curve. **D**) FVB/NJ growth ratio exceeds balanced growth curve r. **E**) C57Bl/6J L6-7 folding index shows limited folding increase. **F**) FVB/NJ L6-7 folding index reveals increase in folding. For all analysis C57Bl/6J = 38; FVB/NJ = 41.

### Cerebellar strains have similar geometry and growth ratio at folding initiation

Previously we provided evidence that cerebellar folding emerges without a molecular prepattern determining the location of the folds (Lawton et al., 2019). Still, it could be that differences in the initial size or geometry of the cerebella of each strain, could give rise to the differences in the amount of folding that become apparent later in development. However, we found that at E16.5 the strains have the same midsagittal cerebellar area, EGL length, and geometric relationship between their length and area (Supplemental Fig 3F-H). We also compared the growth ratio of the EGL and core at the initiation of folding (from E16.5 until the cerebella reach a size of 1mm^2^, corresponding to ∼P0 for both strains). Multiple linear regression analysis showed that there was no difference between the EGL/core growth ratios (Supplemental Fig 3I-L). Together these results provide evidence that the difference in folding amount is not pre-figured by changes in the geometry of the cerebellar anlagen or embryonic growth ratios.

### The growth ratio is increased in L6-7 of FVB/NJ compared to C57Bl/6J

We next tested whether the EGL/core growth ratio was different between the strains within the L6-7 region where the level of folding diverges. Given that the anchoring centers maintain fixed positions within the cerebella and that growth within each lobule region is constrained within the lobules, it is possible to determine regional levels of expansion and folding by treating the anchoring centers as boundaries (Lawton et al., 2019; Legué et al., 2015; Sudarov and Joyner, 2007; Szulc et al., 2015). We measured the expansion rate of L6-7 starting at the beginning of the critical period (roughly FVB/NJ: P3, C57Bl/6J: P5) when the area of L6-7 was at least 0.3mm^2^, until the area was no more than 0.8mm^2^ (roughly P6 for FVB/NJ and P8 for C57Bl/6J). Multiple linear regression analysis showed that the growth ratio (EGL length/cerebellar area) of L6-7 of C57Bl/6J cerebella is reduced when compared with FVB/NJ (Fig. 3A).

### The level of differential-expansion is regionally distinct and correlates with the cerebellar folding amount

We next asked if the growth ratios (EGL/core) are regionally regulated within each strain. We first measured the growth ratio of lobules 4-5 (L4-5) and lobule 8 (L8) from the same individuals as the measurements for L6-7. In C57Bl/6J both L4-5 and L8 had reduced EGL/core growth ratios when compared with their FVB/NJ counterparts. (Supplemental Fig 4A,B). Multiple Linear regression analysis showed that in C57Bl/6J there was no difference between the EGL/core growth ratios of L4-5, L6-7, or L8. Within FVB/NJ, L6-7 is statistically different than L4-5 though the change is minute and may not be biologically relevant (Supplemental Fig 4C-E).

A differential-expansion between layers emerges when the EGL/core growth ratio exceeds a balanced expansion defined by the tissue geometry (Supplemental Fig 4F). To determine if the measured difference in growth ratios between the strains constitutes a change in the level of EGL/core differential-expansion we constructed balanced-expansion curves from the lobule regions by isometrically scaling the median lobule region from the start of the critical period (L6-7 regions with an area of 0.3 - 0.4 mm^2^, Fig. 3A,B). If the growth ratio is maintained along the balanced-expansion curve, then the growth is balanced for the geometry and no folding will occur. However, a ratio of growth where the slope exceeds the curve indicates the presence of differential-expansion with the EGL growing faster than the core and predicts folding.

We found that L6-7 in C57Bl/6J has less EGL/core differential-expansion than L6-7 in FVB/NJ through the critical period (Fig 3 C,D). The C57Bl/6J strain remains centered around the balanced growth line for longer, and when the rate of EGL expansion increases above the curve it remains closer to the curve compared to FVB/NJ. The difference in the level of EGL/core differential-expansion is also visualized in the residuals of fitting the data to the predictive balanced-expansion curve (Supplemental Fig. 4G). Correspondingly, both the timing of folding, and amount of folding follow the onset and degree of EGL/core differential expansion in both strains (Fig. 3 E,F).

While the sizes of L6-7 are the same between the strains at the start of the critical period (Fig 2E,F) the predictive balanced-expansion curves shows that the C57Bl/6J cerebellum requires a higher level of length expansion to produce a balanced expansion than that required in FVB/NJ (Fig 3 C,E). This indicates that there is a difference in the geometry of L6-7 between the strains. Indeed, plotting the lobule length against the area revealed that L6-7 in C57Bl/6J has a lower area to length ratio than FVB/NJ (Supplemental Fig 4H,I). This shift in the ratio results in the requirement for a steeper EGL/core growth ratio to induce the same level of differential-expansion as in FVB/NJ. Therefore, by adjusting both the EGL/core growth ratios and the tissue geometry, a cerebellum can set the level of EGL/core differential-expansion and control the amount of folding in L6-7.

Within FVB/NJ the three neighboring lobule regions, L4-5, L6-7 and L8, all have very similar EGL/core growth ratios (Supplemental Fig 4C). However, these lobule regions achieve different folding amounts during the critical period with L8 remaining unfolded, L4-5 folding once, and L6-7 folding twice. Therefore, we postulated that the distinct geometry of each lobule region must be defining unique balanced expansion ratios that must be overcome for the individual lobule regions to fold. We focused on Lobule 8 because of its simpler shape. In both strains lobule 8 is largely constrained by the surrounding lobule regions. As a result, its width does not increase as the lobule grows. This non-isometric expansion can be compared with a rectangle that has a fixed width and a growing length (Supplemental Fig 5A,B). For this type of growth, the predicted balanced-expansion profile takes a linear form (Supplemental Fig 5C). We found that in both strains the growth ratio of L8 closely approximates the geometrically determined balanced-expansion profile during the critical period, and as predicted, produces no folding (Supplemental Fig 5D,E). Therefore, while the growth ratios between L4-5, L6-7, and L8 are similar within each strain the resulting level of differential-expansion within each strain is regionally regulated by the distinct tissue geometries of the individual lobule regions. Lastly, the small increase in the folding index observed in L4-5 in C57Bl/6J during the critical period comes from the complex geometry and not the onset of new fissures (Supplemental Fig 5F,G).

### The EGL thickness regionally varies between strains

In bi-layer models of cerebral cortex folding, the thickness of the outer layer has been predicted to regulate the wavelength of the resulting folds with thicker layers predicted to produce greater wavelengths between folds (Tallinen et al., 2014). To test if the thickness of the EGL during the formation of the anchoring centers (base of the fissure) could control the folding wavelength, we measured the thickness of the EGL during the critical period when the first fissure that subdivides L6-7 appears.

We measured the thickness of the EGL by lobule region at the start and end of the critical period (P5, P7 for C57Bl/6J and P3, P5 for FVB/NJ). The EGL thickness varied across the lobule regions in both strains (Figure 4A-C, Supplemental Fig 6A). At the start of the critical period, prior to the formation of the anchoring center, the EGL of L6-7, L8 and L9 is thicker in C57Bl/6J than in FVB/NJ. At the end of the critical period, thickness of the EGL in L6, L8, and L9 are unchanged between the strains while the EGL of L7 in C57Bl/6J is thicker compared with FVB/NJ (Supplemental Fig 6A). At the end of the critical period the collective L6-7 is not statistically different between the strains (data not shown).

**Fig 4.**
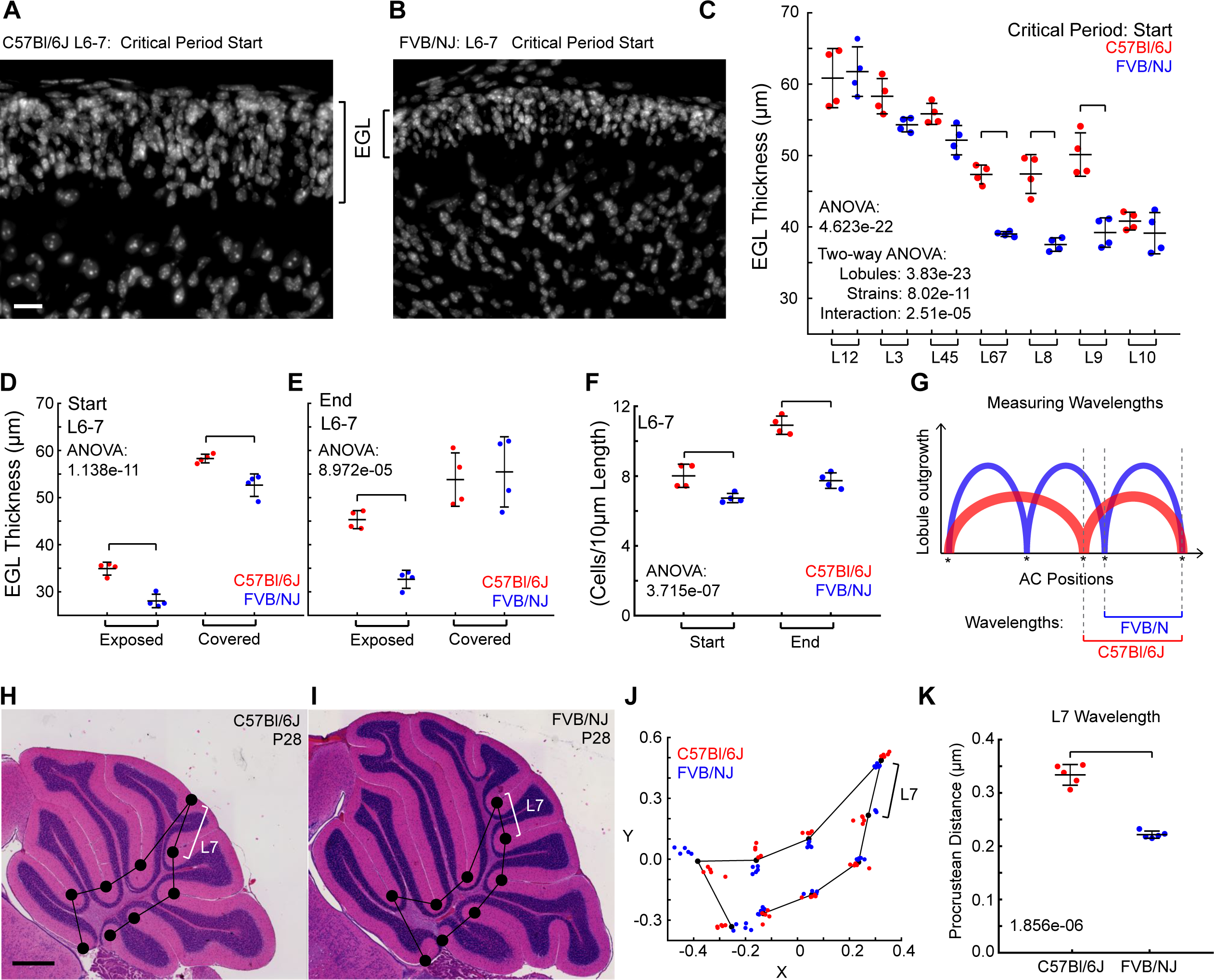
The EGL thickness correlates with the wavelength. **A-B**) EGL of L6-7 region. Scalebar: 20µm **C-F** n = 4/strain **C**) EGL thickness is regionally regulated within and between the strains. **D-E**) Exposed vs covered EGL thickness in L6-7 during the critical period. **F**) Number of cells per 10µm of L6-7 EGL length. **G)** Cartoon of wavelength measurements. **H,I**) Cerebella with placement of the landmarks at the conserved anchoring centers. White Brackets indicate L7 wavelength. Scale bar: 500µm **J**) Landmark-based procrustean analysis. n = 5/strain. Bracket indicates L7 wavelength **K**) Wavelength of Lobule 7 is increased in C57Bl/6J cerebella (n = 5/strain). (Mean, s.d.)

We next tested if the EGL thickness varied within the regions of L6-7, those that are exposed (where the fissures will appear) and those that are covered (adjacent to other lobules) (white and cyan in Supplemental 6B). At the start of the critical period the EGL of the exposed region was thinner in both strains than the covered region. However, the difference between the strains was most pronounced in the exposed region with the C57Bl/6J having a thicker EGL than FVB/NJ at the start and end of the critical period. By the end of the critical period the EGL in the covered regions showed no difference in thickness between the strains (Fig. 4D,E). As another way to measure the thickness of the exposed surface of L6-7 we quantified the number of cells per EGL surface length (Fig 4F). As expected, the C57Bl/6J cerebella had more cells per EGL length in L6-7 than in FVB/NJ, and the density of the EGL between the strains was the same (Supplemental Fig 6C).

### Final Wavelength of Folding in L6-7 is predicted by EGL thickness

To test if the thicker EGL in C57Bl/6J results in an increased folding wavelength we measured the direct distance between the anchoring centers shared by both strains at P28 (Fig.4 G-I). Anchoring centers (ACs) largely hold their positions in space and therefore retain the spatial information of the EGL surface from the period when they were placed (Sudarov and Joyner, 2007; Szulc et al., 2015). Further, each AC has a robustly stereotyped timing of appearance (Legué et al., 2015). Therefore, the thickness of the EGL at the time of each AC formation should contribute to the final wavelength of the enclosed lobule. Since the entire C57Bl/6J cerebellum is only ∼72-75% the size of FVB/NJ we used a landmark-based procrustean analysis to correct for the global size difference (Fig 4J).

Within each strain, each of the ACs showed tight clustering with minimal variation (Fig. 4J and Supplemental Fig 6D-F). Using the positions of the ACs retained in both strains, we measured the wavelength of each lobule as the direct distance between the two ACs on either side of each lobule region. The wavelength of L7 was increased in C57Bl/6J compared to FVB/NJ as predicted while the wavelengths of L1-2 and L3 were decreased in C57Bl/6J. (Fig 4K, Supplemental Figure 6G). The remaining wavelengths were unchanged between the strains even L8 and L9 whose EGL thickness was increased at the start of the critical period. However, only the anterior AC enclosing L7 is generated during the critical period when the EGL thickness is increased in C57Bl/6. The anchoring centers setting the boundaries of the other lobules are formed prior to this critical period.

An unchanged lobule wavelength predicts that at the time the ACs formed, the EGL in that region had a similar thickness between the strains. Therefore, we chose to measure the EGL thickness of L8 as its wavelength was unchanged between the strains. The Secondary fissure, emerging embryonically (∼E18), marks the posterior boundary of L8. The Prepyramidal fissure, marking its anterior limit, is already established in the undivided L6-7-8 region at P1 in FVB/NJ and P2 in C57Bl/6J. Therefore, we measured the EGL thickness of L6-7-8 one day prior (P0 and P1 in FVB/NJ and C57Bl/6J, respectively). The EGL thickness was similar between the strains as predicted; however, the small difference was statistically significant suggesting the EGL in FVB/NJ is ∼2µm thicker than in C57Bl/6J. (Supplemental Figure 6I).

We also measured the folding wavelength in absolute distance, not correcting for size differences, and observed the same pattern of wavelengths across the cerebella (Supplemental Fig 6H). All the measured anchoring centers, save the one marking the anterior boundary of L7, are in place prior to the critical period, before any significant differences in the growth ratios of the cerebellum develop (Fig. 2A) and are maintained in position during development.

### Strains have different densities of Purkinje cells at the completion of folding

Since the level of differential expansion and EGL thickness are different between the strains and regionally regulated within the cerebellum, we next sought to determine the cellular mechanism accounting for these differences. We first investigated the cellular players that drive expansion of the EGL and the cerebellum. Purkinje cells provide the mitogen that drives the expansion of GCPs within the EGL through their proliferation (Lawton et al., 2023). Without this cell expansion the folding is severely diminished (Corrales et al., 2006). Purkinje cells also play an important role in scaling other cell populations (interneurons and astrocytes) in the cerebellar cortex to form the appropriate number of cellular partners for functional circuits (Joyner and Bayin, 2022; Willett et al., 2019). Therefore, we investigated if the density of Purkinje cells was different between the strains in a way that could change the level of EGL expansion and therefore the growth ratio, differential expansion, and folding.

We first determined the total count and density of Purkinje cells per midline section in each lobule at P28 (Supplemental Fig. 7A-E). The lobules have unique numbers of Purkinje cells, and L1-2 and L6, had fewer Purkinje cells in C57Bl/6J than in FVB/NJ accounting for the small global reduction in Purkinje cells in C57Bl/6J. However, FVB/NJ was found to have a lower density of Purkinje cells (number/length) than C57Bl/6J at P28. Several lobule regions (L4-5, L6, L9 and L10) showed reduced density in FVB/N. Thus, in regions where the number of Purkinje cells are the same, C57Bl/6J cerebella expand less per Purkinje cell than FVB/NJ. Additionally, in the L6 region where folding is most reduced, the number of Purkinje cells is also reduced in C57Bl/6J.

### During the critical period L6-7 of FVB/NJ has a higher density of Purkinje Cells

We next measured the density of Purkinje cells during the critical period in the L6-7 region (Fig. 5A-D). At the start of the critical period the density of Purkinje cells was reduced in L6-7 of C57Bl/6J mice compared with FVB/NJ. However, by the end of the critical period, after the folding amount has diverged, the density of Purkinje cells equalized between the strains (Fig. 5E). A similar pattern was observed in L4-5 and L8 (Supplemental Fig. 7F,G). In addition, the Purkinje cells in L6-7 were tightly distributed with less distances between nearest neighbors in the FVB/NJ than in C57Bl/6J at the start before reaching the same spatial distribution at the end of the critical period (Fig. 5F).

**Fig 5.**
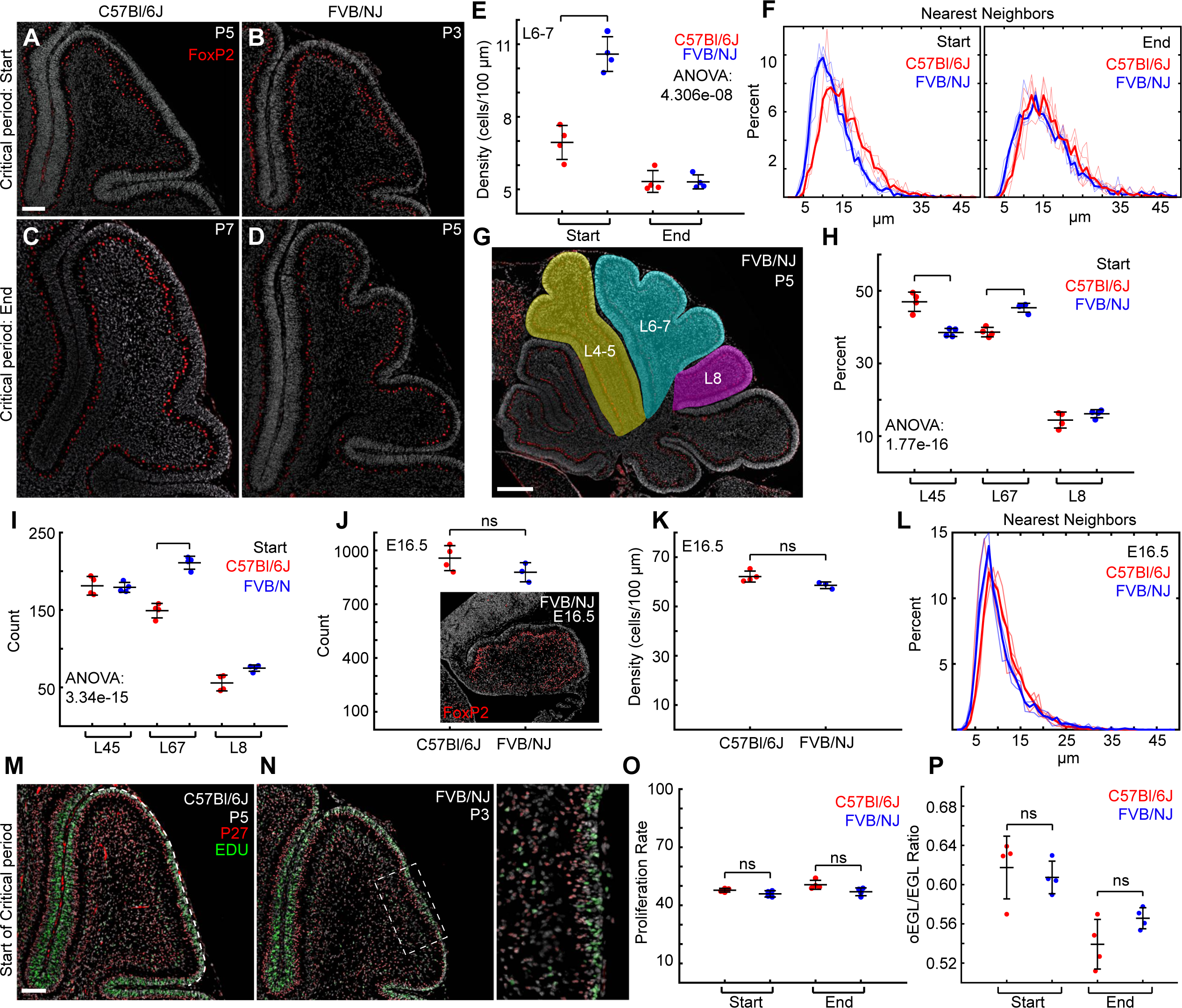
Purkinje cell number predicts folding amount. **A-D**) Sagittal midline sections of L6-7 stained with FoxP2. Scale bar: 100µm **E**) Purkinje Cell density. **F**) Purkinje cell are closely packed in FVB/NJ at the start of the critical period. **G, H**) The percentage of Purkinje cells within L4-5, L6-7, and L8. Scale bar: 300µm **I**). L6-7 has a reduced number of Purkinje cells in C57Bl/6J. **J-L**) Both strains have similar numbers, densities, and distributions of Purkinje cells at E16.5, C57Bl/6J: n = 4, FVB/NJ: n = 3, **M,N**) Sagittal midline sections stained with P27, EDU, and DAPI. Solid line: exposed surface of lobule. Scale bar: 100µm **O**). The proliferation rate is unchanged between strains. **P**) The relative size of the proliferating portion of the EGL is unchanged. (mean, s.d.)

Calculating the density of Purkinje cells at both the start and end of the critical period as a ratio of each lobule’s final density at P28 showed that L6-7 in FVB/NJ is denser throughout the critical period whereas L4-5 and L8 are only higher at the start in FVB/NJ (Supplemental Fig. 7H,I). As the Purkinje cells eventually form a single monolayer throughout the cerebellum, the higher density in FVB/NJ during the critical period of L6-7 could indicate that the regional density of Purkinje cells, and by extension the amount of SHH secreted, may prefigure the final amount of folding through increasing the expansion of the EGL.

### Purkinje cells numbers are reduced in L6-7 during the critical period

We next addressed whether the difference in Purkinje cell density in L6-7 between strains during the critical period was a result of improper cell sorting or differences Purkinje cell number. One possibility is that a portion of the Purkinje cells in C57Bl/6J are improperly directed to the lobules surrounding L6-7. To test this hypothesis, we determined the percent of all Purkinje cells in L4-5, L6-7, and L8 in each strain at the critical period (Fig. 5G,H). In C57Bl/6J a greater percentage of the Purkinje cells were located within L4-5 and the percentage in L6-7 was reduced compared with FVB/NJ (Fig. 5H). Likewise, the number of Purkinje cells was reduced in L6-7 in C57Bl/6J. However, the number of Purkinje cells was the same between the strains in both L4-5 and L8 (Fig. 5I). We next tested if fewer Purkinje cells are present at the start of cerebellum development in C57Bl/6J, specifically those that will occupy L6-7. We measured the numbers of Purkinje cells at the start of folding (E16.5), which is 3 days after Purkinje cell have been born. We found no difference in the number, density, or distribution of Purkinje cells between strains (Fig 5J-L).

### The proliferation rate of granule cells does not regulate the level of differential-expansion

Since the number and density of Purkinje cells is reduced in L6-7 in C57Bl/6J compared to FVB/NJ we hypothesized that the proliferation rate of GCPs within the EGL would be reduced, potentially lowering the level of differential-expansion. We measured the proliferation rate [EDU+/P27-] of the GCPs in the exposed surface of L6-7 at the start and end of the critical period (Fig 5M,N). Surprisingly, we found no differences in the rate of proliferation at the start or end of the critical period between strains (Fig 5O).

The proliferation of the EGL is constrained to the outer P27-cells of the EGL (oEGL) (green in Fig 5M,N). We hypothesized that the oEGL may be larger in FVB/NJ, containing more cells able to proliferate, and leading to more expansion with the same rate of proliferation. However, the oEGL/EGL ratio is the same between the strains (Fig. 5P). These results indicate that cell proliferation within the EGL is not adjusted to set the growth ratios or the resulting level of differential-expansion in L6-7 between the strains.

### Cell division angle predicts EGL tangential expansion and thickness

Cell division angle (CDA) within the EGL corresponds to the bias in EGL expansion in the anterior-posterior direction as opposed to the medial-lateral direction during cerebellum development (Legué et al., 2015). Additionally, it was found that removing CHD7 from GCPs affects their division angle – increasing the proportion of cells that divide medial-laterally (Reddy et al., 2021). Under this change the cerebellum showed lobule like folds in a medial-lateral pattern. Therefore, we hypothesized that CDA could be a fundamental mechanism to regulate both the level of differential-expansion and the thickness of the EGL, ultimately controlling both the folding amount and folding wavelength.

We measured the CDA in L4-5, L6-7 and L8 by labeling for Phospho-Histone H3 and determining the division plane in reference to the surface of the cerebellum (Fig 6 A,B). We predicted to find a biases towards vertical divisions (60-90 degrees from the surface) in FVB/NJ given the increased tangential expansion and reduced thickness of its EGL. Excitingly, we found that throughout the critical period, FVB/NJ has a higher proportion of vertical divisions in L6-7 compared with C57Bl/6J (Fig. 6C-G, Supplemental Fig 8A-C). At the start of the critical period, collectively, FVB/NJ is biased towards vertical divisions (supplemental Fig 8E) compared with C57Bl/6J. But by the end of the critical period the bias towards vertical divisions in FVB/NJ was only maintained in L6-7 (Fig 6G, Supplemental Fig 8A-C).

**Fig 6.**
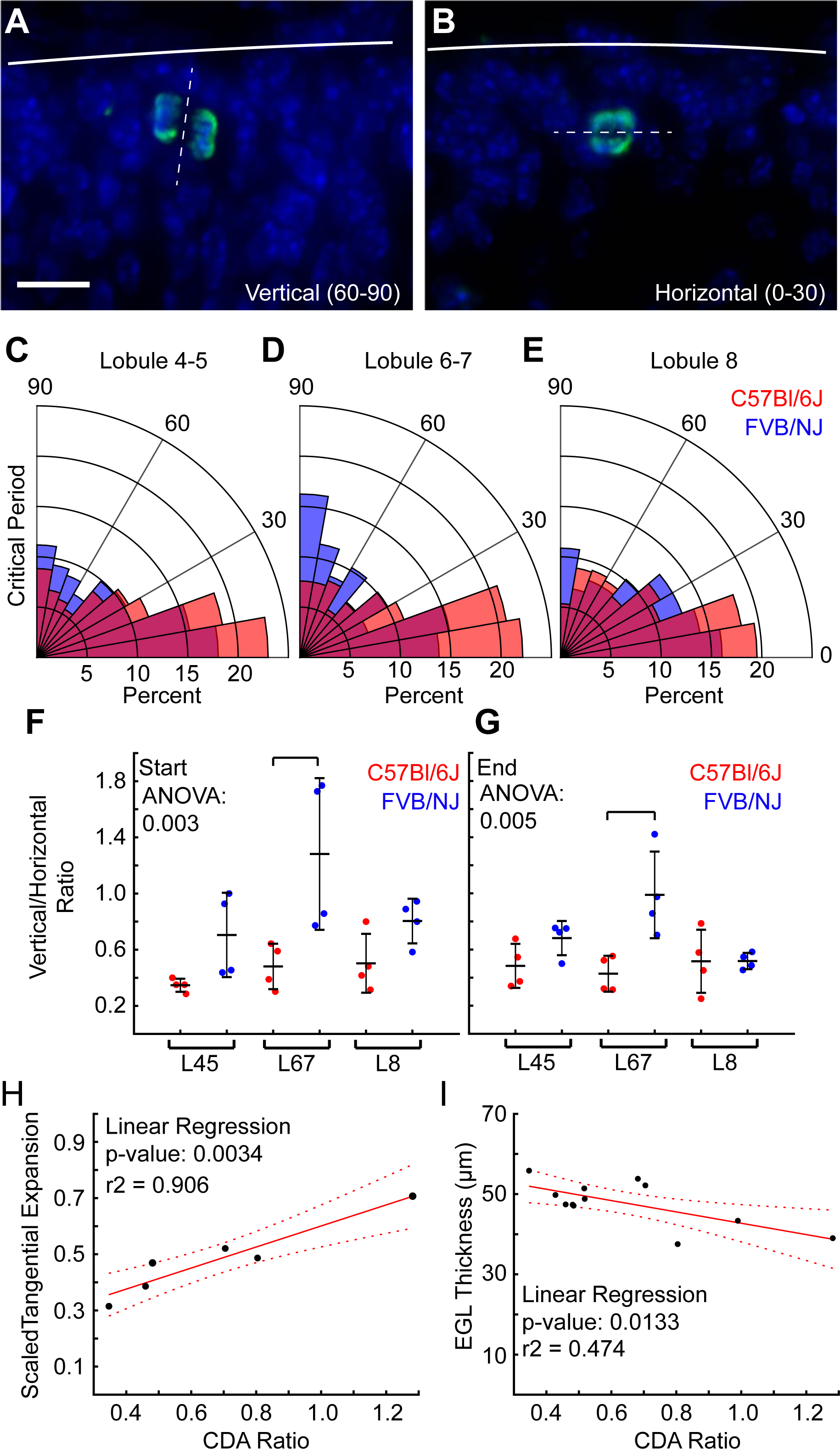
Cell division angle is altered between strains. **A,B**). PH3 staining showing vertical and horizontal divisions within the EGL. Scale bar: 10µm **C-E**) Rose plots of CDA in L4-5, L6-7, and L8. n = 8/strain. **F,G**) Cell division ratio is biased vertically in FVB/NJ compared to C57Bl/6J in L6-7, n = 4/strain. **H,I**) Cell division ratio predicts tangential expansion and EGL thickness. (mean s.d.)

We next tested the relationship of CDA to the tangential expansion of the EGL. We measured the average tangential expansion of the EGL in L4-5, L6-7, and L8 in the cerebella used for the cell division measurements during the critical period. We then ran linear regression analysis using the cell division ratio for each lobule region to predict the tangential expansion of the EGL during the critical period (Fig 6H). We found a significant fit with a high r-squared value demonstrating that the CDA explains over 90% of the variation in the EGL tangential expansion. Similarly, we tested the relationship of the CDA to the EGL thickness. Again, we found a statistically significant relationship between the division angle and the EGL thickness. However, the CDA only explains some 47% of the variability. This indicates that the CDA is only one factor, of potentially several, that contribute to the regulation of EGL thickness (Fig 6I).

## Discussion

Here we tested multiple predictions for neural tissue folding during cerebellar foliation. We found that the degree of cerebellar folding correlates with regional levels of differential-expansion due to changes in both the underlying growth ratio between the EGL and the core and to the geometry of the lobules. We also provided developmental evidence that the folding wavelength is regulated by adjusting the thickness of the EGL at the time of fissure formation. We further propose that the angle of cell division within the EGL is a tunable regulator that affects both the tangential expansion and thickness of the EGL (Fig 7 A-C).

**Fig 7.**
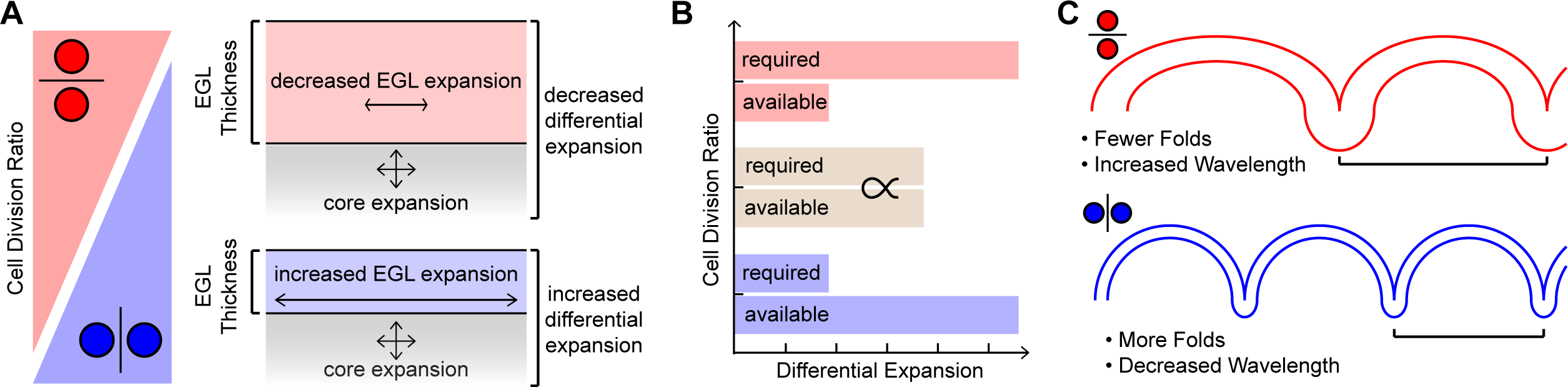
Cell Division Angle regulates the level of folding by modulating the tangential expansion of the EGL and its thickness. **A**) High vertical cell division ratios increase the tangential expansion and decrease the thickness of the EGL. Low vertical ratios increase the thickness of the EGL while decreasing the tangential expansion. **B**) Increasing the thickness of the EGL and decreasing its tangential expansion increases the force required to fold the tissue and decreases the force available in the tissue. **C**) Cell division angle regulates folding amount and wavelength.

### Gene and time-agnostic approaches allow for studies of cerebellar intrinsic mechanics

The gene-agnostic approach allowed an unbiased analysis of the tissue mechanics predicted to regulate the degree of cerebellar folding. We propose that our approach of comparing inbred mouse strains provides an approximation of the natural folding variation seen between individuals within a species. Intriguingly, the folding differences studied here are similar to those seen across healthy human populations where global variation is minimal and larger changes in amount are constrained to specific regions (Luders et al., 2006, 2004; Zilles et al., 1988). An alternative approach to ours to capture the mechanical regulation of natural folding variation would be to use an outbred strain that has substantially more variation between individuals than inbred strains. But without the folding robustness seen in C57Bl/6J and FVB/NJ, a direct comparison of different individuals at different developmental timepoints would be impossible.

The time-agnostic analysis of cerebellar growth rate (Fig 2A) allowed us to identify a critical period during development, intrinsic to each strain, when the tissue mechanics diverged. Given that the clock-based rate of development runs slower in C57Bl/6J than in FVB/NJ mice, a dependence of clock-time would have reduced the power of the comparisons between the strains. The fact that FVBN/J reaches the same size and degree of folding, ∼2 days prior to C57Bl/6, illustrates that the folding amount is intrinsic to the tissue mechanics of the cerebellum and not the global speed of development.

### The level of Differential-Expansion is regionally adjusted to set the folding amount

We uncovered differences in the lobule EGL/core growth ratios and resulting levels of differential expansion between FVB/NJ and C57Bl/6J. First, the tangential expansion of the EGL was increased in FVB/NJ compared to in C57Bl/6J in all lobules measured. The differences in L6-7 between the strains was further magnified by the slight geometric difference in the starting shape of L6-7 in each strain. This shape change determines the required ratio of EGL to core growth needed to achieve the same differential-expansion required to lead to folding. In both strains L4-5 and L8 showed very similar growth ratios to L6-7. Yet the divergent geometries significantly modulate the resulting level of differential-expansion needed to produce additional folding. Our data indicates that cerebella can regionally altered both the EGL/core growth ratio and the lobule geometry to adjust the level of cerebellar folding.

### EGL thickness at the time of fissure formation is correlated with the folding wavelength

The thickness of the EGL during the critical period was varied within and between the strains, as well as within individual lobules. Excitingly, the significant greater thickness of L6-7 in C57Bl/6 compared to FVB/NJ, at the time when the fissures surrounding L7 form, correlates with a longer wavelength of the resulting lobule. In contrast, L8, which showed no difference in the wavelength between strains at P28 had a very similar EGL thickness at the time of fissure formation. The slight difference in thickness detected is a fraction of a single cell diameter and is likely not biologically significant.

### Purkinje cell density during the critical period predicts folding amount

Given the role of GCPs in the growth, expansion, and folding of the cerebellum, we predicted that the level of proliferation would be a tunable regulator of the EGL to core growth ratio and therefore of differential-expansion of layers and folding (Corrales et al., 2006; Legué et al., 2016). Surprisingly, our data argues that the proliferation rate within the EGL of L6-7, is not tuned to adjust the level tangential expansion of the EGL (Fig 5O).

In contrast, we found a regional reduction in the number and density of Purkinje cells in L6-7 of C57Bl/6J at the start of the critical period correlating with the future degree of folding (Fig 5E,I). Given that the cerebella of the two strains have the same number of total Purkinje cells in the midline at E16.5 and that the lobules adjacent to L6-7 have comparable numbers of Purkinje cells as their FVB/NJ counterparts during the critical period, we hypothesis that the Purkinje cells in the L6-7 region of C57Bl/6J undergo increased cell death or move a greater distance medial-laterally compared with FVB/NJ mice.

Surprisingly, Purkinje cell densities at P28 were elevated in C57Bl/6J compared with FVB/NJ. This shows that FVB/NJ achieves more growth per Purkinje cell than C57Bl/6J and suggests that there may be strain intrinsic differences in the degree to which Purkinje cells are able to drive the expansion of other cell populations to scale the proper balance of cells within the cerebellum (Joyner and Bayin, 2022; Willett et al., 2019). Indeed, there are many fixed mutational differences between C57Bl/6J and FVB/NJ mice, and indeed, variations also between sub-strains. (Bowes et al., 1990; Kasugai et al., 2007; Mekada and Yoshiki, 2021; Schauwecker, 2012; Timmermans et al., 2017). Together, our findings suggests that both the regional number of Purkinje cells during development, as well as their strain intrinsic effectiveness could have a role in scaling the folding of the circuitry.

### Cell division angle as a tunable regulator of the tangential expansion and thickness of the EGL

In *Lkb1* mutant mice cerebellar folding is increased and EGL thickness is decreased without changing the proliferation rate in the EGL, but with an increase in vertical cell divisions (Ryan et al., 2017). Additionally, lobule-like structures form in the mutant cerebellum in the medial-lateral direction when the cell division angle is biased in that direction (Reddy et al., 2021). By shifting the CDA the cerebellum may adjust various mechanical parameters that regulate the degree and wavelength of folding. A horizontal bias should increase the thickness of the EGL at the expense of its tangential expansion. The increased thickness should increase the force required to fold; simultaneously the reduction in tangential expansion will reduce the level of EGL to core differential-expansion, the driving force for folding, and reduce folding (Fig 7A-C).

Supporting this model, the angle of cell division in the EGL is biased vertically in L6-7 of FVB/NJ compared to C57Bl/6J precisely when we measured a reduced EGL thickness and greater tangential-expansion of the EGL compared to in C57Bl/6 (Fig. 6C-G). Further, at the start of the critical period in FVB/NJ, there is an increased bias to vertical divisions compared to C57Bl/6J and each lobule region in FVB/NJ has a higher tangential expansion than in C57Bl/6J (Fig 3A, Supplemental Fig 4A,B1, 8E). Similarly, at the end of the critical period when L4-5 and L8 have similar division ratios within and between the strains, the resulting EGL thicknesses within and between the strains are similar (Supplemental Fig. 6A, 8A,C).

While we detected a statistically significant relationship between the angle of cell division and the EGL thickness, it only accounts for 47% of the observed variation in EGL thickness. This result predicts that other factors work together with the division angel to tune the EGL thickness. It is plausible that the migration mechanics of GCPs, or their rates of differentiation could play a role. Or it may be that tensile forces, predicted to cross the EGL, may also affect the EGL thickness (Lawton et al., 2019). Excitingly, the cell division angle predicts over 90% of the variation in the tangential expansion of the EGL, suggesting it is a dominant regulator of the tangential expansion and by extension, folding amount of the cerebellum.

### Materials and Methods Animals

Animals were maintained in accordance protocols approved by the Institutional Animal Care and Use Committees at Mississippi State University and Memorial Sloan Kettering Cancer Center. All data was collected from the two inbred mouse strains, C57Bl/6J (Jackson Labs: 000664) and FVB/NJ (Jackson Labs: 001800). Both sexes were used for analysis. Mice were maintained on a 12hr light/dark cycle and food and water were provided ad libitum.

The appearance of a vaginal plug was used to mark noon as Embryonic day 0.5. Pups were injected subcutaneously with 25ug/g 5-ethynyl-2deoxyruidine (EdU; Invitrogen) one hour prior to collection.

### Tissue Preparation and Imaging

#### Adult Stage

P28 cerebella dissected out of the skull at noon and fixed in Bouin’s solution at room temperature and rotated for 24 hours. Brains were then washed in PBS, dehydrated, and prepared for paraffin embedding. Parasagittal sections were collected with a Leica Microtome RM2235 at 8µM.

#### Developmental Series

All cerebella of developmental stages were collected at approximately noon on the day of collection. Post-natal brains were dissected out of the skull and postfixed in fresh, ice-cold 4% PFA for 24 hours. Embryonic heads were postfixed as above. Brains were washed in PBS and cryoprotected with washes of 15% sucrose and 30% sucrose in PBS. Brains were frozen in OCT, stored at −80C and cut with a Leica CryoStat (CM3050s) for 10µM parasagittal sections.

The slides with the most midline sections were chosen and stained with H&E. All sections were Imaged on a Zeiss Observer Z.1 with Apatome, or Leica Thunder Imaging System.

### Area, Length, Positive curvature, Folding Index measurements, and fissure counts

Measurements were collected either in Imaris (Bitplane) or ImageJ as previously described (Lawton et al., 2019). Three midline sections were measured from each cerebellum and the median values were reported. The global length was measured from the anterior start of the surface of the Molecular layer (P28) or the anterior start of the EGL (developmental series). The area was measured by joining the anterior and posterior ends by following the ventricular zone. In ImageJ the positive curvature was created using the convex hull to delete all negative curvature points. Regional, lobule-based measurements were similarly collected. Lobule lengths, Positive curvature and folding were measured from the surface of the molecular layer/EGL at the anchoring center to the anterior of the lobule to the center of the anchoring center posterior to the lobule region. Lobule region areas were measured by started at the base of the EGL directly below the surface of the AC anterior to the lobule region of interest. The area then followed the surface of the EGL until it reached the posterior anchoring center enclosing the lobule region. The area was extended directly below the surface of the AC to the bottom of the EGL. Then the area was directly closed by connecting to the start.

For the adult stage five brains were measured per strain. For the developmental series, C57Bl/6J were collected daily from E16.5 to P9 and FVB/NJ were collected daily from E16.5 to P7. For the developmental series 94 cerebella were measured for C57Bl/6J and 96 for FVB/NJ. For each brain in the series the 2-3 most midline sections were measured, and the median values were reported.

### Antibodies and EdU staining

Antibody and EdU staining was performed as previously described (Lawton et al., 2019). Prior to IHC EdU was detected using a commercial kit (Invitrogen C10340). Primary antibodies were incubated overnight at 4C or at Room temperature for 4 hours: Goat anti-Foxp2 Everest (1:1000, EB05226), rabbit anti-PH3 EMD Millipore (1:1000, 06-570), Mouse anti-P27 BD Biosciences (1:500, 610241), rabbit anti-Calbindin Swant (1:500, CB38). All antibodies were diluted in 2% milk and 0.2% Triton X-100. AlexaFluor secondary antibodies at a 1:1000 dilution were used and sections were counter stained with DAPI.

### Cell Counts, Proliferation, EGL Thickness, Division Angle

Purkinje cells were counted in Imaris. Four cerebella were used per strain and per timepoint for developmental stages. For each brain 6-9 midline sections were measured. At P28 five brains were used and four midline sections were measured per brain. The median value for each brain was reported. For E16.5 Measurements four brains were used for C57Bl/6J and three for FVB/NJ and 3 sections per brain were measured. The nearest neighbor analysis was calculated in Imaris software. Matlab (Mathworks) software was used to remove the duplication bias that arose when a pair of cells is nearest neighbors with themselves.

The Proliferation rate was measured as previously described (Lawton et al., 2019). Four cerebella were used per each strain and per time point. Three midline sections per brain were measured. The median proliferation rate for each brain was reported. All cells within the EGL of the exposed portion of the lobule were measured. DAPI+ cells that were both EDU+ and P27-were counted as proliferating. The proliferation rate was calculated as (number of EDU+ cells) / (number of DAPI+; P27-cells).

The thickness of the EGL was first measured by dividing the area of the EGL by its length. In the L6-7 region the thickness was also measured in the exposed and covered portions of the EGL. Lastly the thickness and density of the exposed region of L6-7 was measured by counting all of the cells within the EGL dividing the counts by the length of the exposed surface.

The cell division angle was measured in Imaris software. Four brains were measured per strain and per timepoint. For each brain, 6-9 midline sections were measured. The median values per brain were reported. The angle of cell division was measured in relation to the local surface of the EGL.

To test the relationship between cell division angle and EGL tangential expansion the cell division angle was measured in L4-5, L6-7 and L8 in four brains at the start of the critical period and the average of the four brains was taken for L4-5, L6-7, and L8. The EGL length was also measured in those brains as well as the four brains at the end of the critical period. The average length difference between the start and end for each lobule region was normalized to control for the differences in their starting sizes. The 6 average cell division ratios (L4-5, L6-7, L8 from C57Bl/6J and FVB/NJ) from the start of the critical period were plotted against the normalized EGL expansion during the critical period. Similarly, the average cell division ration for each lobule region at the start and end of the critical period was plotted against the average EGL thickness measured in the same brains at the start and end.

### Size-free wavelength measurements

We used MorphoJ software to run a standard land-mark based procrustean analysis to control for the size difference between the strains (Klingenberg, 2011). We used the 9 conserved anchoring centers as the landmarks and ran standard alignments individually and with strains together.

### Statistics

All statistics analyses were run using Matlab (Mathworks). When individual comparisons were made the two-sample t-test was used. The statistical threshold was a p<0.05. When multiple comparisons were being made one-way ANOVA analyses were performed. After ANOVA analysis a multiple comparison was run with Tukey’s honestly significant difference criterion. The statistical threshold was set at a p< 0.05. When testing for differences between the strains, the lobule regions, and any interactions, two-way ANOVA analyses was performed and the p-values for strain, lobule, and interaction were reported. All error bars are standard deviations. No statistical methods were used to predetermine the sample sizes. We used sample sizes aligned with the standards in the field. No randomization was used nor was data collected or analyzed blind. See statistics table.

### Regression Analyses

For the global growth curve fitting we ran non-linear regressions using a basic form of the Gompertz function:

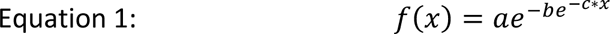

Initial coefficients for the equation were estimated through repeated graphical exploration and Excel solver to minimize residuals.

Linear regression analysis was done in Matlab to test for fits. Multiple linear regression analysis was done to test for differences between slopes. See Statistics Table

## Acknowledgements

We would like to thank the members of the Lawton and Joyner labs for their discussion and support. We thank Dr Nathan Wisnoski for discussion on statistical analysis.

## Funding

This work was supported by grants from the NSF (CAREER: 2045759) to AKL and NIH (NINDS R01NS092096, NIMH R37MH085726) to ALJ and a National Cancer Institute Cancer Center Support Grant (P30 CA008748-48).

## Data Availability

All relevant data is provided in the article and its supplementary files.

## Competing Interests

The authors no competing or financial interests.

---

**Supplemental Figure 1:**
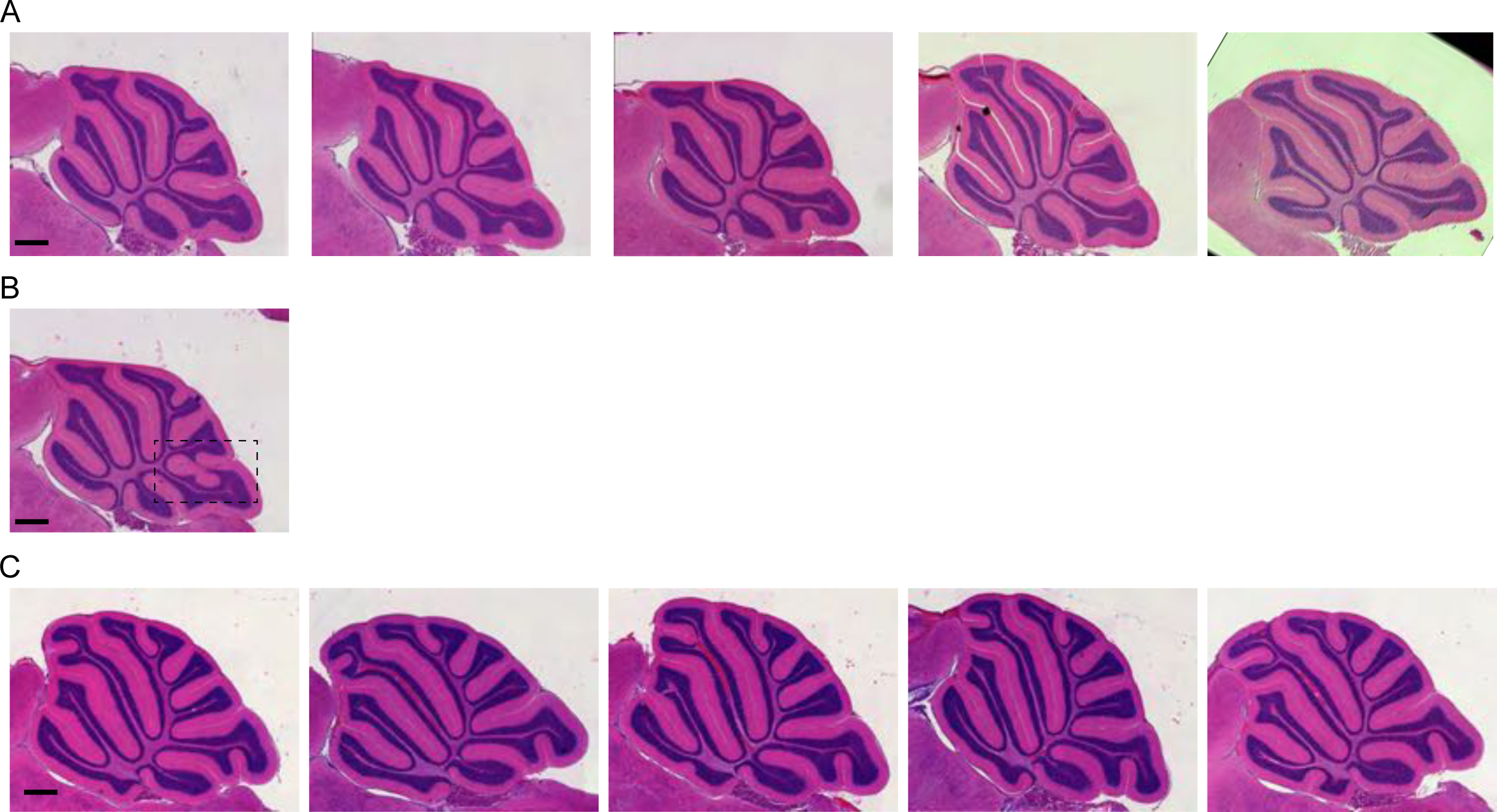
C57Bl/6J and FVB/NJ cerebella at P28 have robustly different levels of folding at the midline of the vermis. **A)** Sagittal midline sections of 5 C57Bl/6J cerebella **B)** C57Bl6/J cerebella showing heterotopia between lobule 8 and Lobule 9. **C)** Sagittal midline sections of 5 FVB/NJ. All cerebella were stained with H&E. Scale Bars: 0.5 mm.

**Supplemental Figure 2:**
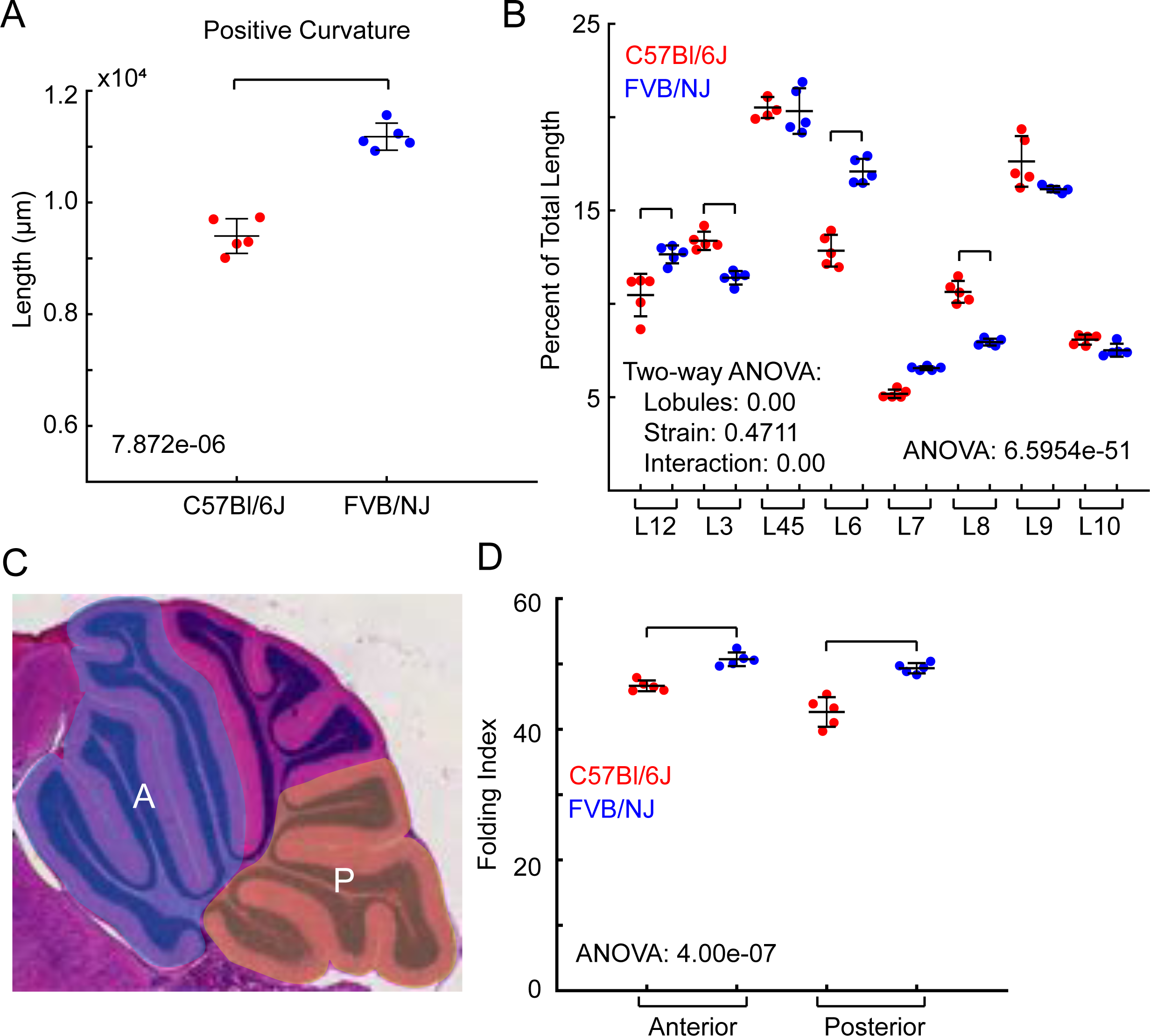
Size difference between C57Bl/6J and FVB/NJ is regionally regulated. **A)** Positive curvature. P-value reported. **B)** Lobule lengths. All regions statistically different (one-way ANOVA p-value reported) except for Lobule 8. Two-way ANOVA (lobules and strains) interaction p-value: 7.68e-20. **C)** Lobule lengths as a percentage of total length. Brackets indicate statistical differences (one-way ANOVA p-value reported). Two-way ANOVA (lobules and strains) interaction p-value: 0.00e+00. **D)** Image showing anterior (blue shading) and posterior (yellow shading) regions of cerebellum. **E)** Folding index of anterior and posterior regions. Brackets indicate statistical differences. One-way ANOVA p-value reported. Two-way ANOVA (Regions and strains) interaction p-value: 0.048. For full statistics see statistics table. (mean s.d.)

**Supplemental Figure 3:**
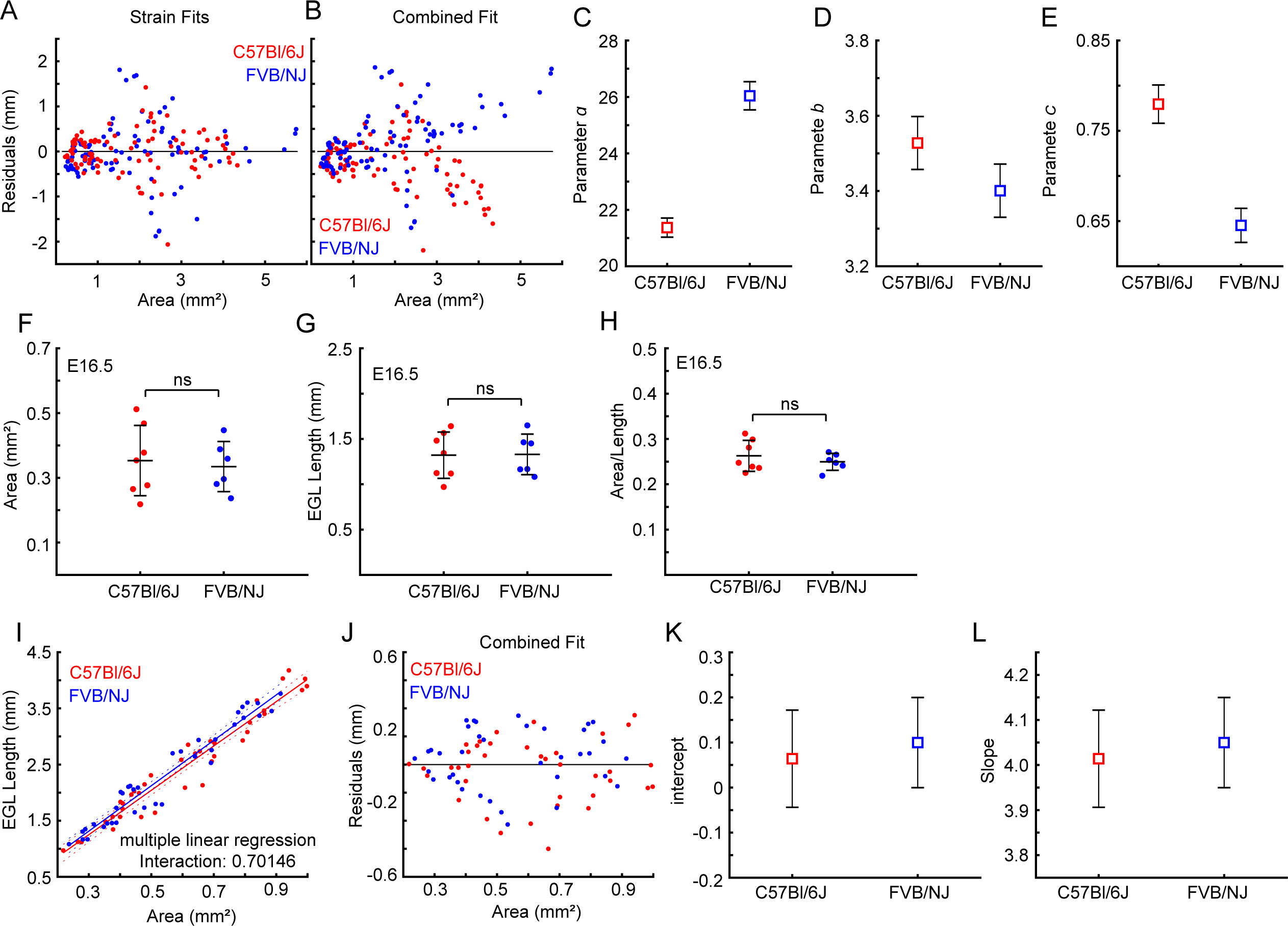
Global ratio of growth diverges during the critical period. **A)** Residuals from fitting each strain to individual Gompertz function (see Fig. 2A). **B)** Residuals are poorly patterned and larger when data from Fig. 2A is combined and fitted to one Gompertz function. **C-E)** The three parameters of the individual Gompertz function fittings are distinct between the strains. **F-H)** At E16.5 the area (p-value 0.7353), length (p-value: 0.9473), and the ratio of the area and length (p-value 0.417) are unchanged between the strains. **I)** Growth ratio from the start (E16.5) to 1mm2 (∼P0). Multiple linear regression analysis interaction p-value reported showing no difference between the slopes. **J)** Residuals are small and well patterned with a single combined fit. **K,L)** The parameters of the individual fittings are overlapping showing no difference between the strains at this early period of growth. For full statistics see statistics table. (mean s.d.)

**Supplemental Figure 4:**
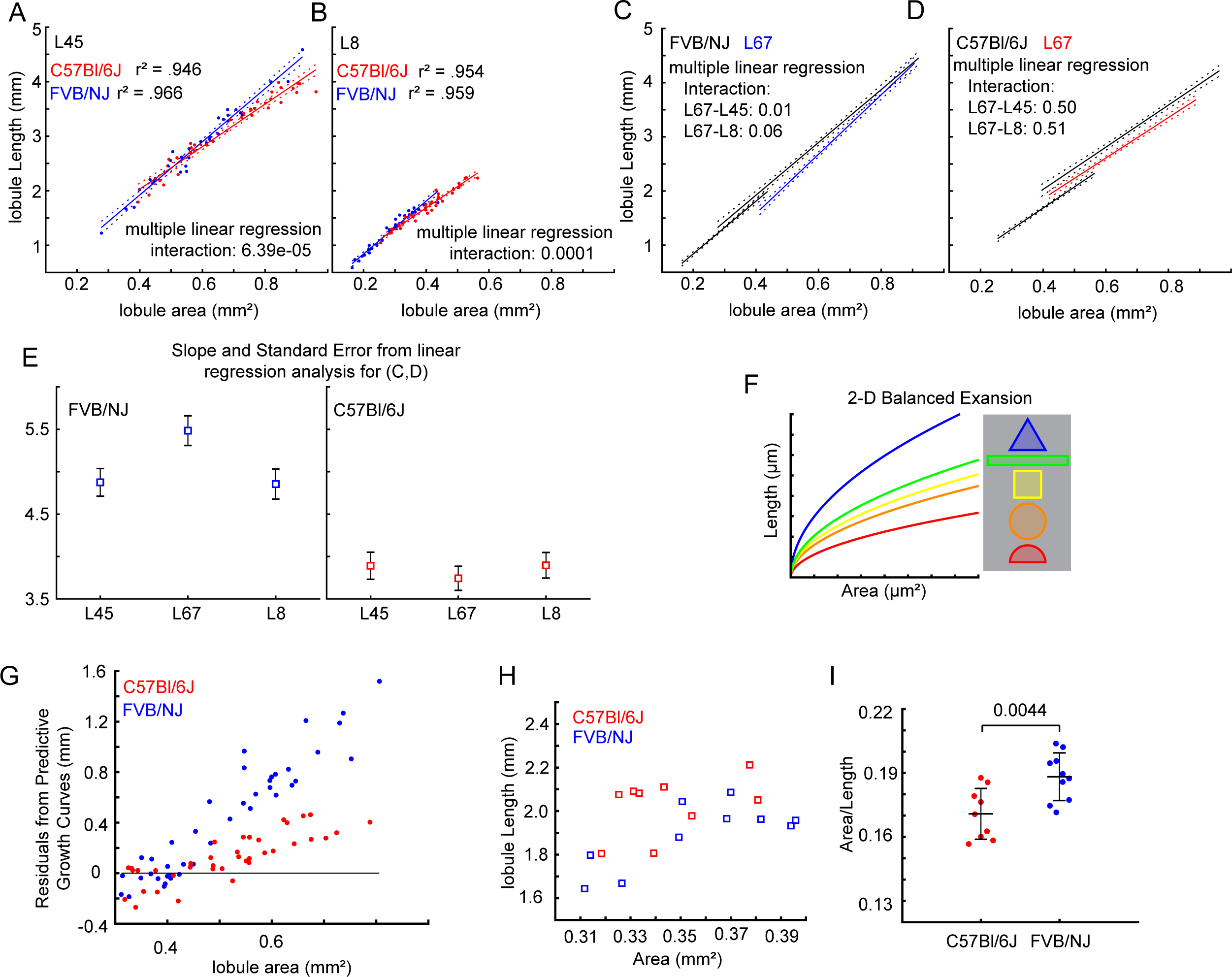
Lobule growth ratios are higher in FVB/NJ than in C57Bl/6J and the resulting differential-expansion is dependent of the geometry of the lobule. **A)** Multiple linear regression analysis of L4-5 region. Interaction p-value reported. Showing difference in slopes. R-squared values reported for individual fits. **B)** Multiple linear regression analysis of L8 region. Interaction p-value reported. Showing difference in slopes. R-squared values reported for individual fits. **C-D)** Multiple linear regression analysis within each stain. Interaction terms for differences between L6-7 and L4-5 or L8 reported. **E)** Calculated slope parameters from linear regression analysis of L4-5, L6-7, and L8 for both strains showing that the slopes are all increased in FVB/NJ compared to C57Bl/6J and the greatest increase is in L6-7. **F)** Cartoon depicting balanced growth ratio (length/Area) curves for common 2-D shapes. **G)** The residuals calculated from the predictive growth curves show that the growth ratio of C57Bl/6J is more similar to its balanced growth curve than FVB/NJ. H,I) L6-7 has a slight difference in geometry between the strains with C57Bl/6J having more length per area than FVB/NJ. P-value reported. (mean s.d.)

**Supplemental Figure 5:**
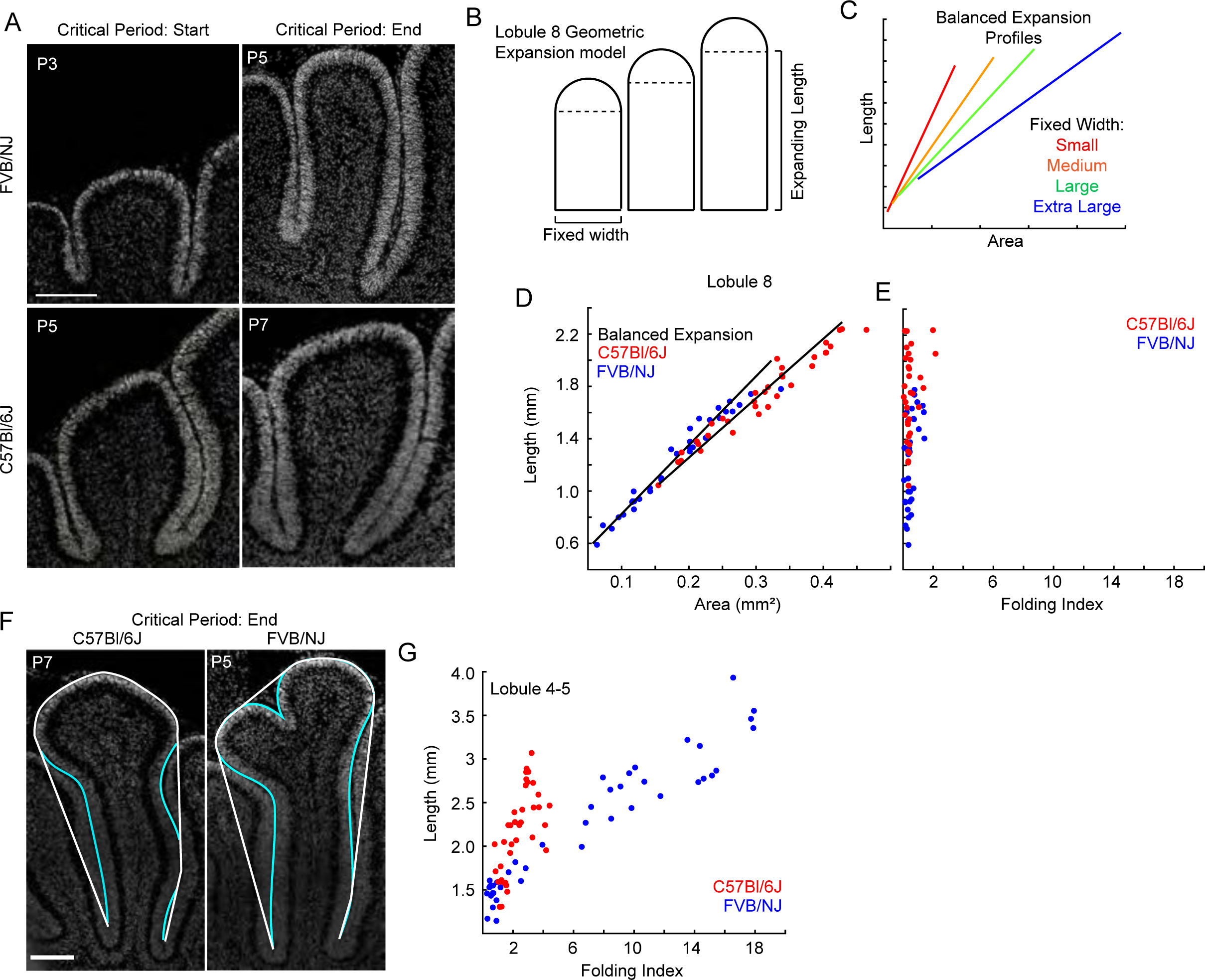
Lobule geometry regionally regulates folding amount. **>A)** Sagittal midline sections of L8 region stained with Dapi at the start and end of the critical period in FVB/NJ and C57Bl/6J showing the constraints from the surrounding lobule regions and the limited exposed surface Scale bar: 200µm **B)** Model of Lobule 8 expansion. The width of the lobule which sets the parameters for the semi-circle is fixed while the length is allowed to expand. **C)** Balanced expansion curves for such constrained shapes are linear and the slope decreases as the fixed width is increased. **D)** The growth ratio of L8 is well predicted in C57Bl/6J and FVB/NJ by this type of constrained growth showing no evidence of differential-expansion. **E)** The folding index shows that L8 in both strains remains unfolded as its growth ratios remain balanced. **F)** Sagittal midline sections of L45 stained with Dapi at the end of the critical period. Cyan line: EGL length. White line: Positive curvature. Scale bar: 200µm **G)** Folding index of L45 through the critical period. While L4-5 in C57Bl/6 remains unfolded the measured increase in folding index comes from the complex shape of the lobule region.

**Supplemental Figure 6:**
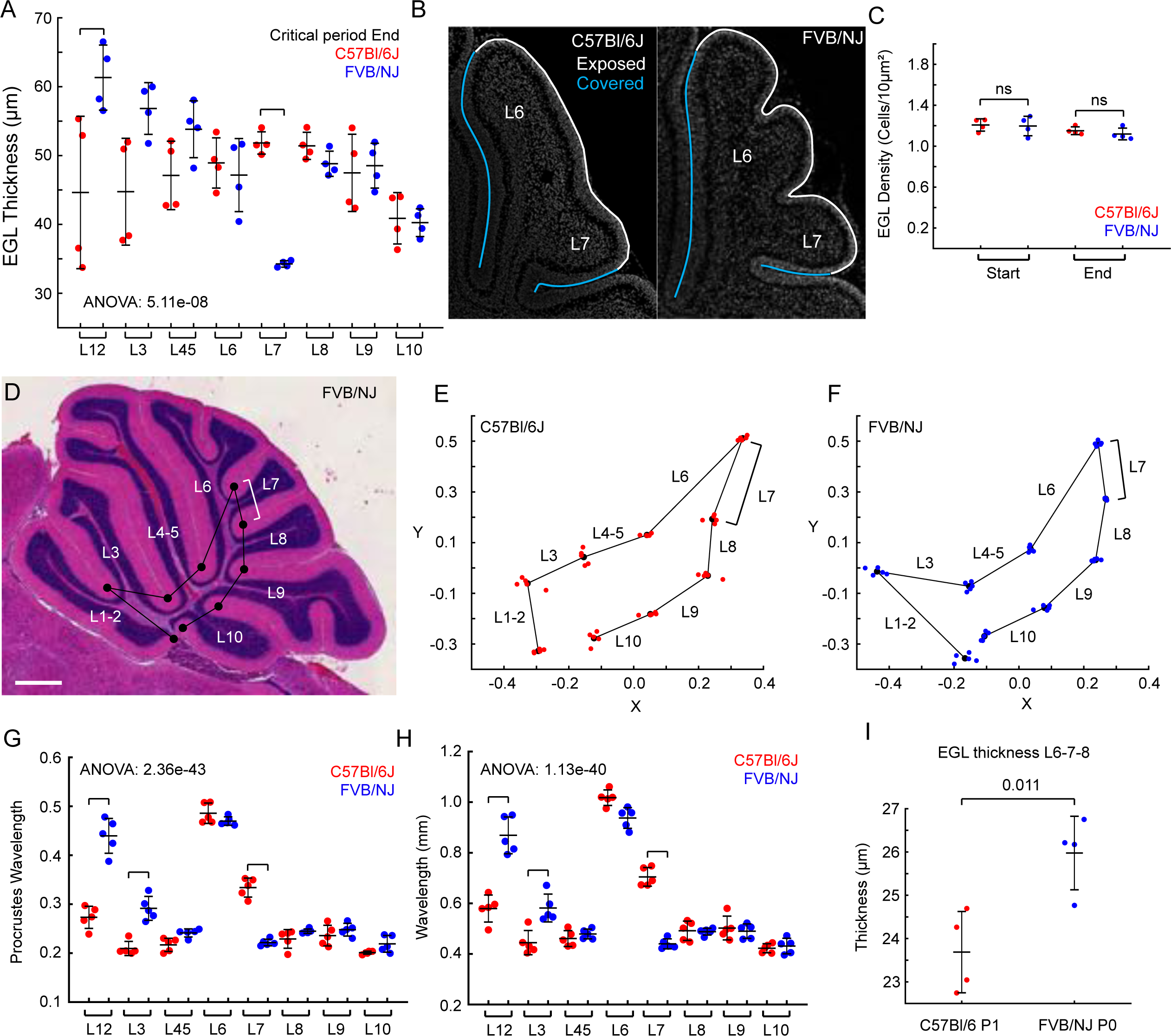
EGL thickness is regionally varied within the cerebellum and correlates with folding wavelength. **A)** EGL thickness at end of critical period. Brackets indicate regions with statistical difference. One-way ANOVA p-value reported. Two-way ANOVA interaction (strain and lobule) p-value: 0.00. **B)** Midline sagittal sections of C57Bl/6J and FVB/NJ L6-7 stained with Dapi. Cyan line: covered EGL. White line: exposed EGL surface. **C)** EGL density in exposed surface. One-way ANOVA p-value: 0.2534. **D)** Midline sagittal section of FVB/NJ at P28 showing landmarks placed at the conserved anchoring centers. Black lines show lobule wavelengths. White bracket indicates L7 wavelength. Scale bar: 0.5mm **E,F)** Individual Procrustes alignments of landmarks of C57Bl/6J and FVB/NJ bracket indicates L7 wavelength. **G)** Wavelengths from Procrustes alignment for each lobule region. Brackets indicate statistical differences. One-way ANOVA p-value reported. Two-way ANOVA interaction (strains and lobules) p-value: 1.71e-23. **H)** Wavelengths from real distances. Brackets indicate statistical differences. One-way ANOVA p-value reported. Two-way ANOVA interaction (strains and lobules) p-value: 0.00 **I)** EGL thickness in L6-7-8 region at P0 and P1 for FVB/NJ and C57Bl/6J. p-value reported. (mean s.d.)

**Supplemental Figure 7:**
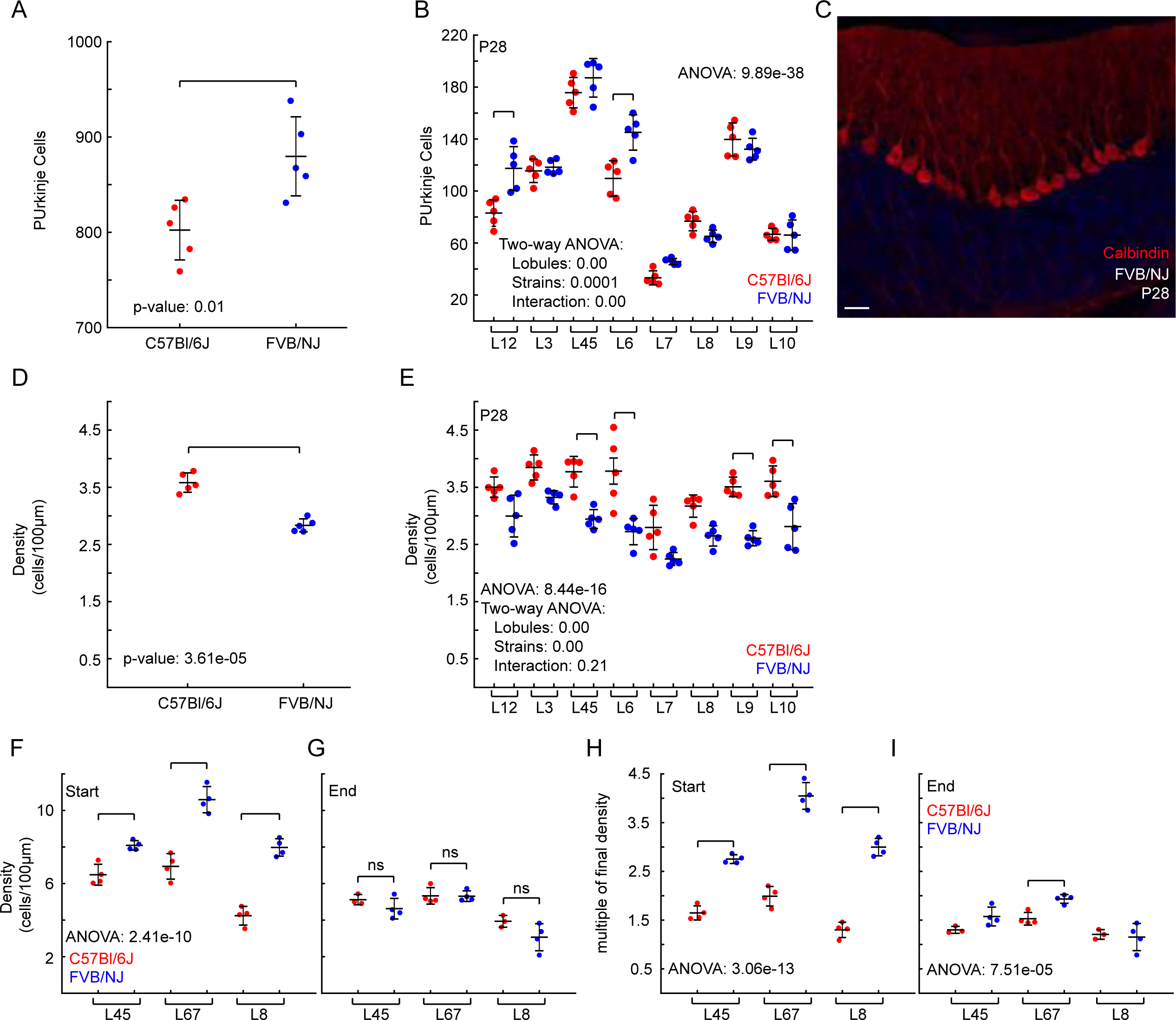
Purkinje cell density is regionally regulated in the Cerebellum and different between the strains. **A)** Number of Purkinje cells per lobule at P28. Brackets indicate statistical differences. One-way ANOVA p-value reported. Two-way ANOVA interaction (strains and lobules) p-value 0.00 **B)** Purkinje cell density at P28. C57Bl/6J has higher density of Purkinje cells even L6 that has a reduced number compared to FVB/NJ. Brackets indicate statistical differences. One-way ANOVA p-value reported. Two-way ANOVA interaction (strains and lobules) p-value: 0.2054 indicating that while both have different densities, they have the same pattern of density. **C)** Sagittal midline section of FVB/NJ at P28 stained with Calbindin and Dapi to mark the Purkinje cells. Scale bar: 50µm **D,E)** Purkinje cell density during the critical period in L4-5, L6-7, and L8. Brackets indicate statistical differences. ns = not statistically significant. One-way ANOVA p-values reported. **F,G)** Purkinje cell density during the critical period as a multiple of final density at P28. Brackets indicate statistical differences. One-way ANOVA p-values reported. (mean s.d.)

**Supplemental Figure 8:**
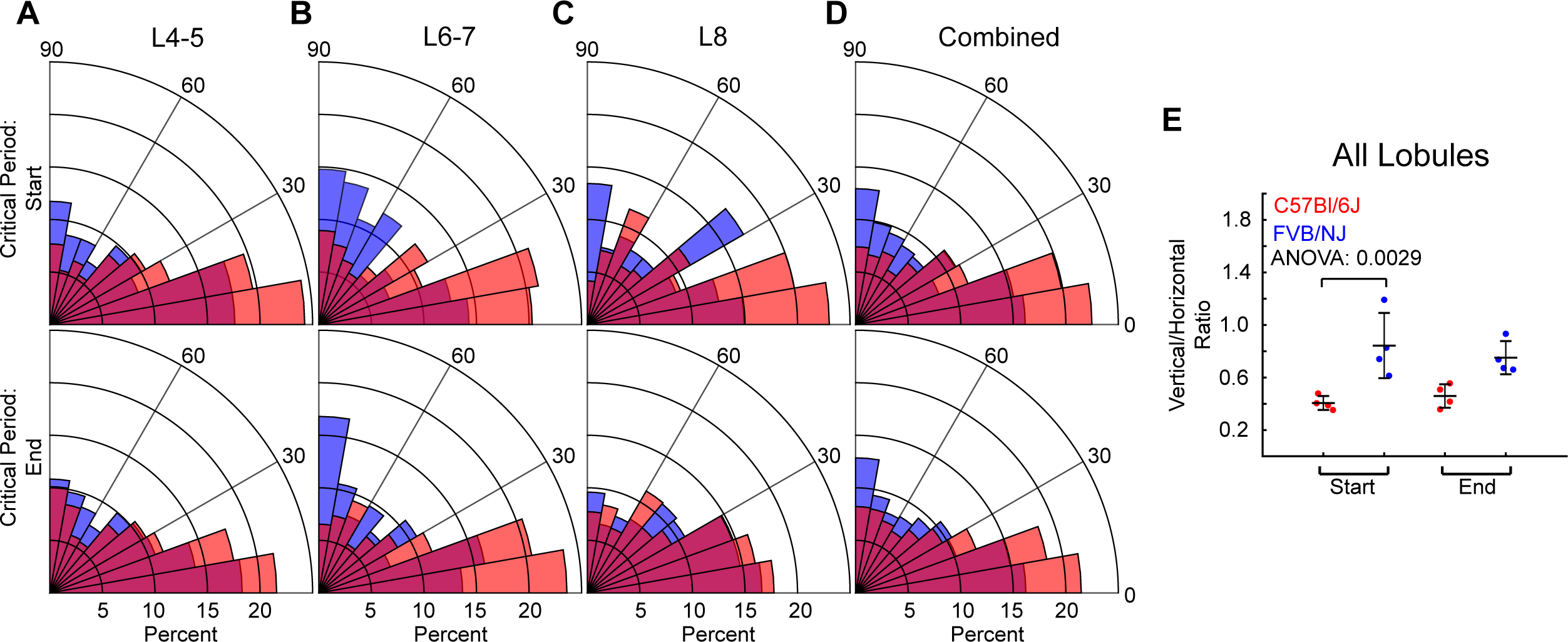
Cell Division Angle is regionally adjusted during the critical period. **A-C)** Polar plot of cell division angles measured in L4-5, L6-7, and L8 at the start and end of the critical periods. At the end of the critical period the difference between the strains is mostly contained to L6-7. **D)** Combined cell division angles measured from L4-5, L6-7, and L8. **E)** Cell division angle ratio of combined data (D). One-way ANOVA p-value reported. Bracket indicates statistical difference. (mean s.d.)

|  |  |  |  |  |
| --- | --- | --- | --- | --- |
| **Figure 1:** |  |  |  |  |
|  | Test | p-value |  |  |
| 1C | ttest2 | 1.83E-05 |  |  |
| 1D | ttest2 | 4.85E-08 |  |  |
|  | Test | p-value | Test | p-value |
| 1E | two-way anova |  | one-way anova | 8.07E-51 |
|  | lobules | 6.96E-51 |  |  |
|  | Strain | 6.05E-34 |  |  |
|  | Interaction | 7.68E-20 |  |  |
| 1F | ttest2 | 7.37E-07 |  |  |
| 1G | ttest2 | 5.23E-09 |  |  |
| **Figure 2:** |  |  |  |  |
|  | Test | p-value |  |  |
| 2A | *Non-linear Regression |  |  |  |
| 2E | ttest2 | 0.3759 |  |  |
| 2F | ttest2 | 0.0653 |  |  |
| **Figure 3:** |  |  |  |  |
|  | Test | p-value |  |  |
| 3A | *Linear Regression |  |  |  |
| **Figure 4:** |  |  |  |  |
|  | Test | p-value | Test | p-value |
| 4C | two-way anova |  | one way anova | 4.62E-22 |
|  | lobules | 3.38E-23 |  |  |
|  | Strain | 8.02E-11 |  |  |
|  | Interaction | 2.51E-05 |  |  |
| 4D | Test | p-value | Test | p-value |
|  | two-way anova |  | oneway anova | 1.138E-11 |
|  | lobules | 0.00E+00 |  |  |
|  | Strain | 0.00E+00 |  |  |
|  | Interaction | 0.4698 |  |  |
| 4E | Test | p-value | Test | p-value |
|  | two-way anova |  | oneway anova | 8.972E-05 |
|  | lobules | 0.00E+00 |  |  |
|  | Strain | 0.0429 |  |  |
|  | Interaction | 0.0126 |  |  |
| 4F | Test | p-value |  |  |
|  | one-way anova | 3.715E-07 |  |  |
| 4K | Test | p-value |  |  |
|  | ttest2 | 1.856E-06 |  |  |
| **Figure 5:** |  |  |  |  |
| 5E | Test | p-value |  |  |
|  | one-way anova | 4.306E-08 |  |  |
| 5H | Test | p-value | Test | p-value |
|  | two-way anova |  | one-way anova | 1.77E-16 |
|  | lobules | 0.00E+00 |  |  |
|  | Strain | 0.9944 |  |  |
|  | Interaction | 0.00E+00 |  |  |
| 5I | Test | p-value | Test | p-value |
|  | two-way anova |  | one-way anova | 3.34E-15 |
|  | lobules | 3.77E-16 |  |  |
|  | Strain | 7.79E-07 |  |  |
|  | Interaction | 3.37E-06 |  |  |
| 5J | Test | p-value |  |  |
|  | ttest2 | 0.1665 |  |  |
| 5K | Test | p-value |  |  |
|  | ttest2 | 0.0602 |  |  |
| 5O | Test | p-value |  |  |
|  | one-way anova | 0.0185 |  |  |
| 5P | Test | p-value |  |  |
|  | one-way anova | 0.0012 |  |  |
| **Figure 6** |  |  |  |  |
| 6F | Test | p-value | Test | p-value |
|  | two-way anova |  | one-way anova | 0.0028 |
|  | lobules | 0.0626 |  |  |
|  | Strain | 0.0005 |  |  |
|  | Interaction | 0.1825 |  |  |
| 6G | Test | p-value | Test | p-value |
|  | two-way anova |  | one-way anova | 0.0046 |
|  | lobules | 0.1378 |  |  |
|  | Strain | 0.0035 |  |  |
|  | Interaction | 0.0224 |  |  |
| 6H - I | *Linear Regression |  |  |  |
| **Supplemental Fig 2** |  |  |  |  |
| 2A | Test | p-value |  |  |
|  | ttest2 | 7.872E-06 |  |  |
| 2B | Test | p-value | Test | p-value |
|  | two-way anova |  | one-way anova | 6.60E-51 |
|  | lobules | 0 |  |  |
|  | Strain | 0.4711 |  |  |
|  | Interaction | 0 |  |  |
| 2D | Test | p-value | Test | p-value |
|  | two-way anova |  | one-way anova | 4.00E-07 |
|  | lobules | 0.0005 |  |  |
|  | Strain | 0 |  |  |
|  | Interaction | 0.048 |  |  |
| **Supplemental Fig 3** |  |  |  |  |
| 3F | Test | p-value |  |  |
|  | ttest2 | 0.7353 |  |  |
| 3G | Test | p-value |  |  |
|  | ttest2 | 0.9473 |  |  |
| 3H | Test | p-value |  |  |
|  | ttest2 | 0.417 |  |  |
| 3I | Test |  |  |  |
|  | *Linear Regressions |  |  |  |
| **Supplemental Fig 4** |  |  |  |  |
| 4E | Test | p-value |  |  |
|  | ttest2 | 0.0044 |  |  |
| 4F-I | *Linear Regressions |  |  |  |
| **Supplemental Fig 6** |  |  |  |  |
| 6A | Test | p-value | Test | p-value |
|  | two-way anova |  | one-way anova | 5.11E-08 |
|  | lobules | 0.00E+00 |  |  |
|  | Strain | 0.1547 |  |  |
|  | Interaction | 0.00E+00 |  |  |
| 6C | Test | p-value |  |  |
|  | one-way anova | 0.2534 |  |  |
| 6G | Test | p-value | Test | p-value |
|  | two-way anova |  | oneway | 2.36E-43 |
|  | lobules | 3.30E-44 |  |  |
|  | Strain | 1.06E-07 |  |  |
|  | Interaction | 1.71E-23 |  |  |
| 6H | Test | p-value | Test | p-value |
|  | two-way anova |  | oneway | 1.13E-40 |
|  | lobules | 0 |  |  |
|  | Strain | 0.1982 |  |  |
|  | Interaction | 0 |  |  |
| 6I | Test | p-value |  |  |
|  | ttest2 | 0.0111 |  |  |
| **Supplemental Fig 7** |  |  |  |  |
| 7A | Test | p-value |  |  |
|  | ttest2 | 0.010 |  |  |
| 7B | Test | p-value | Test | p-value |
|  | two-way anova |  | one-way anova | 9.89E-38 |
|  | lobules | 0 |  |  |
|  | Strain | 0.0001 |  |  |
|  | Interaction | 0 |  |  |
| 7D | Test | p-value |  |  |
|  | ttest2 | 3.61E-05 |  |  |
| 7E | Test | p-value | Test | p-value |
|  | two-way anova |  | one-way anova | 8.44E-16 |
|  | lobules | 0 |  |  |
|  | Strain | 0 |  |  |
|  | Interaction | 0.2054 |  |  |
| 7F | Test | p-value | Test | p-value |
|  | two-way anova |  | one-way anova | 2.41E-10 |
|  | lobules | 0 |  |  |
|  | Strain | 0 |  |  |
|  | Interaction | 0.0017 |  |  |
| 7G | Test | p-value | Test | p-value |
|  | two-way anova |  | one-way anova | 3.52E-05 |
|  | lobules | 0 |  |  |
|  | Strain | 0.0444 |  |  |
|  | Interaction | 0.2704 |  |  |
| 7H | Test | p-value | Test | p-value |
|  | two-way anova |  | one-way anova | 3.06E-13 |
|  | lobules | 0 |  |  |
|  | Strain | 0 |  |  |
|  | Interaction | 0.0002 |  |  |
| 7I | Test | p-value | Test | p-value |
|  | two-way anova |  | one-way anova | 7.51E-05 |
|  | lobules | 0 |  |  |
|  | Strain | 0.0106 |  |  |
|  | Interaction | 0.0495 |  |  |
| **Supplemental Fig8** |  |  |  |  |
| 8E | Test | p-value |  |  |
|  | one-way anova | 0.0029 |  |  |

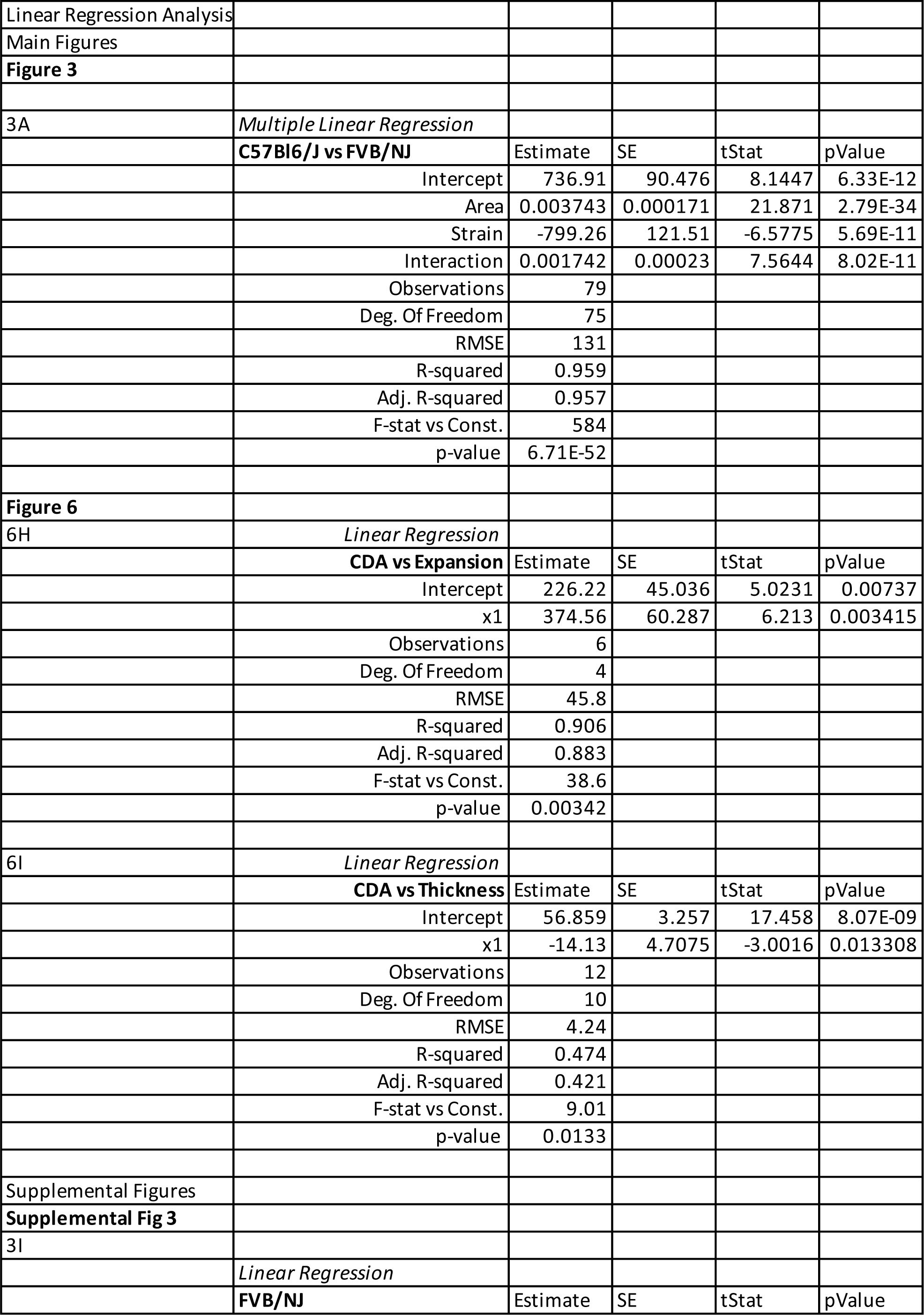
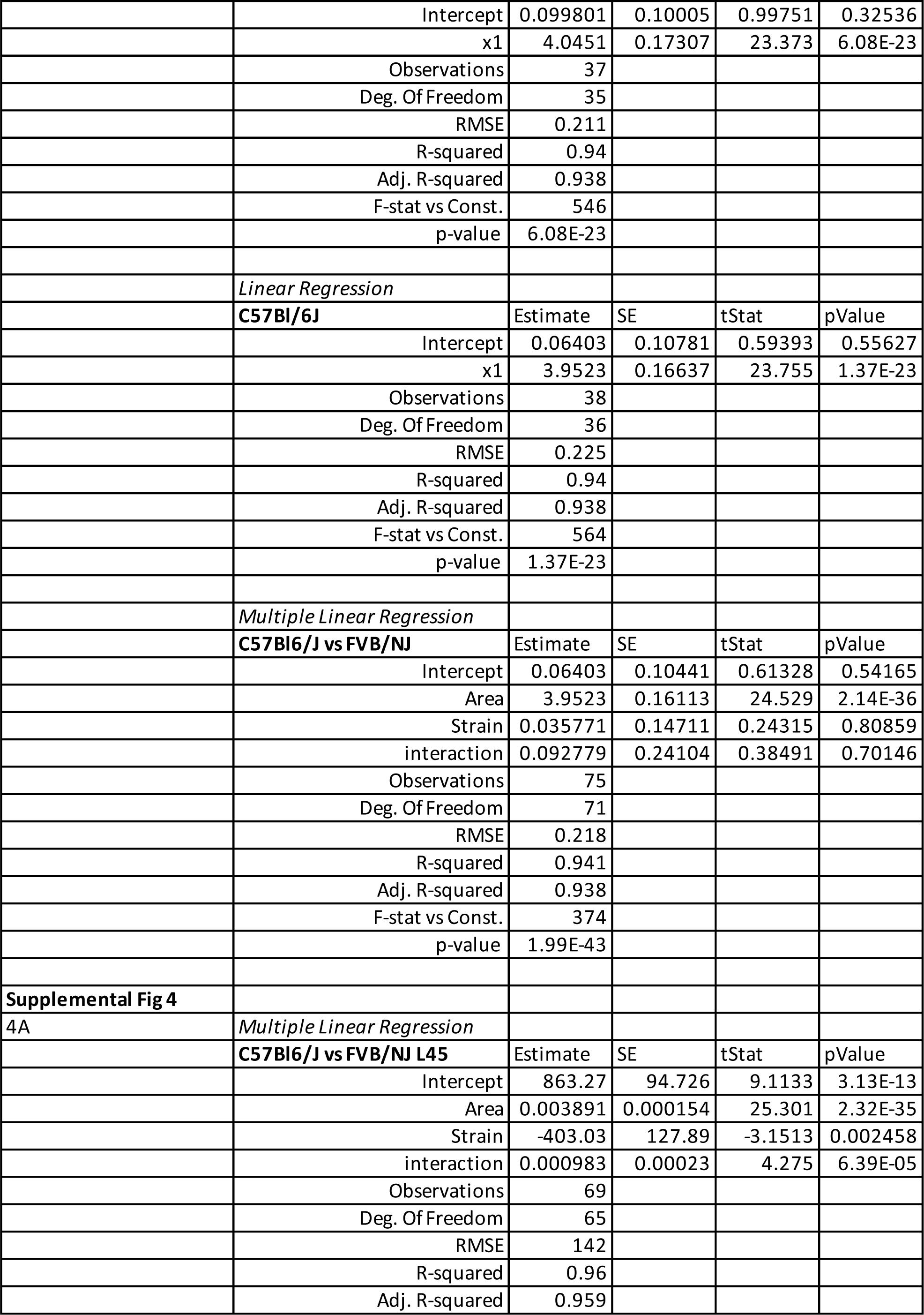
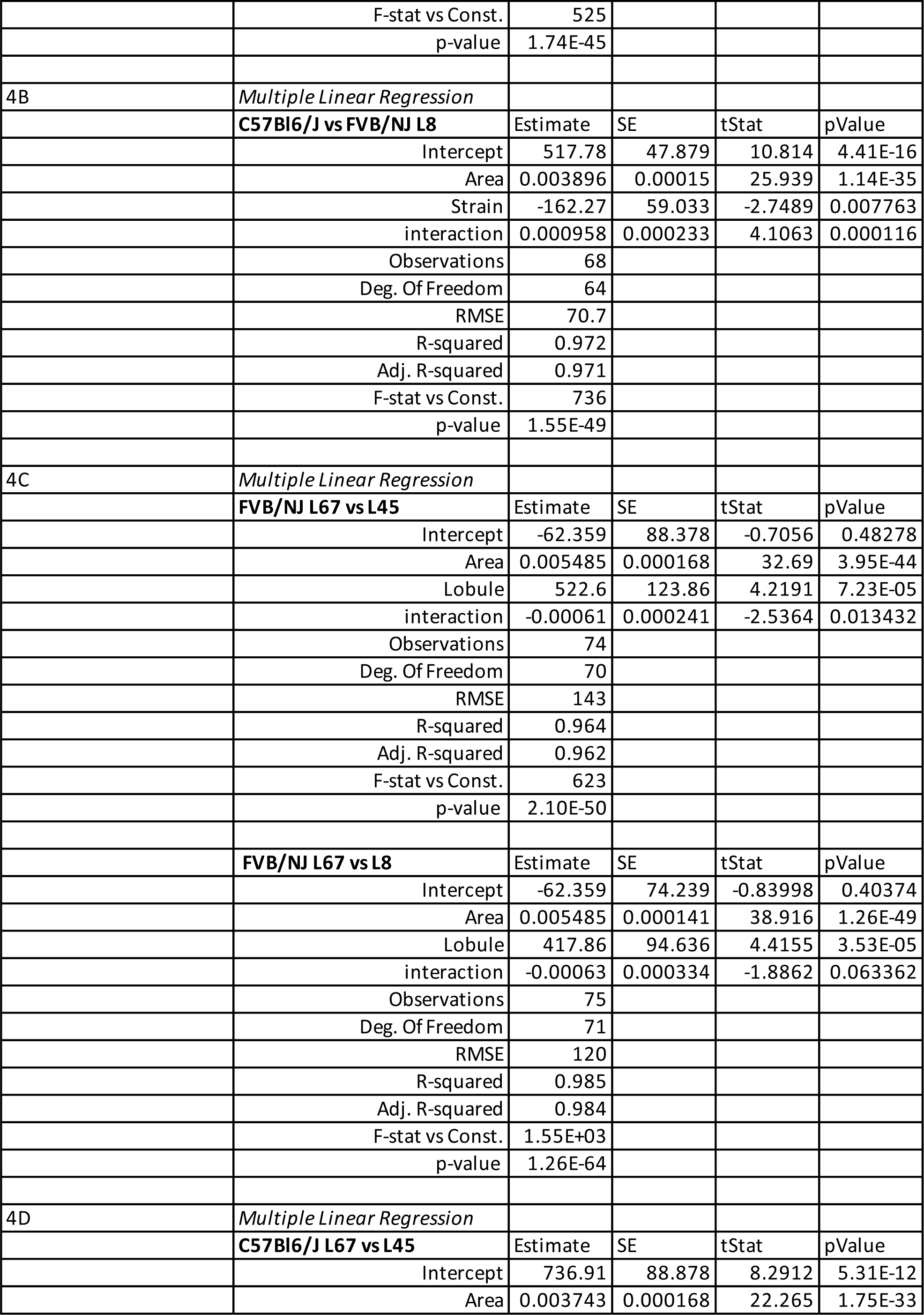
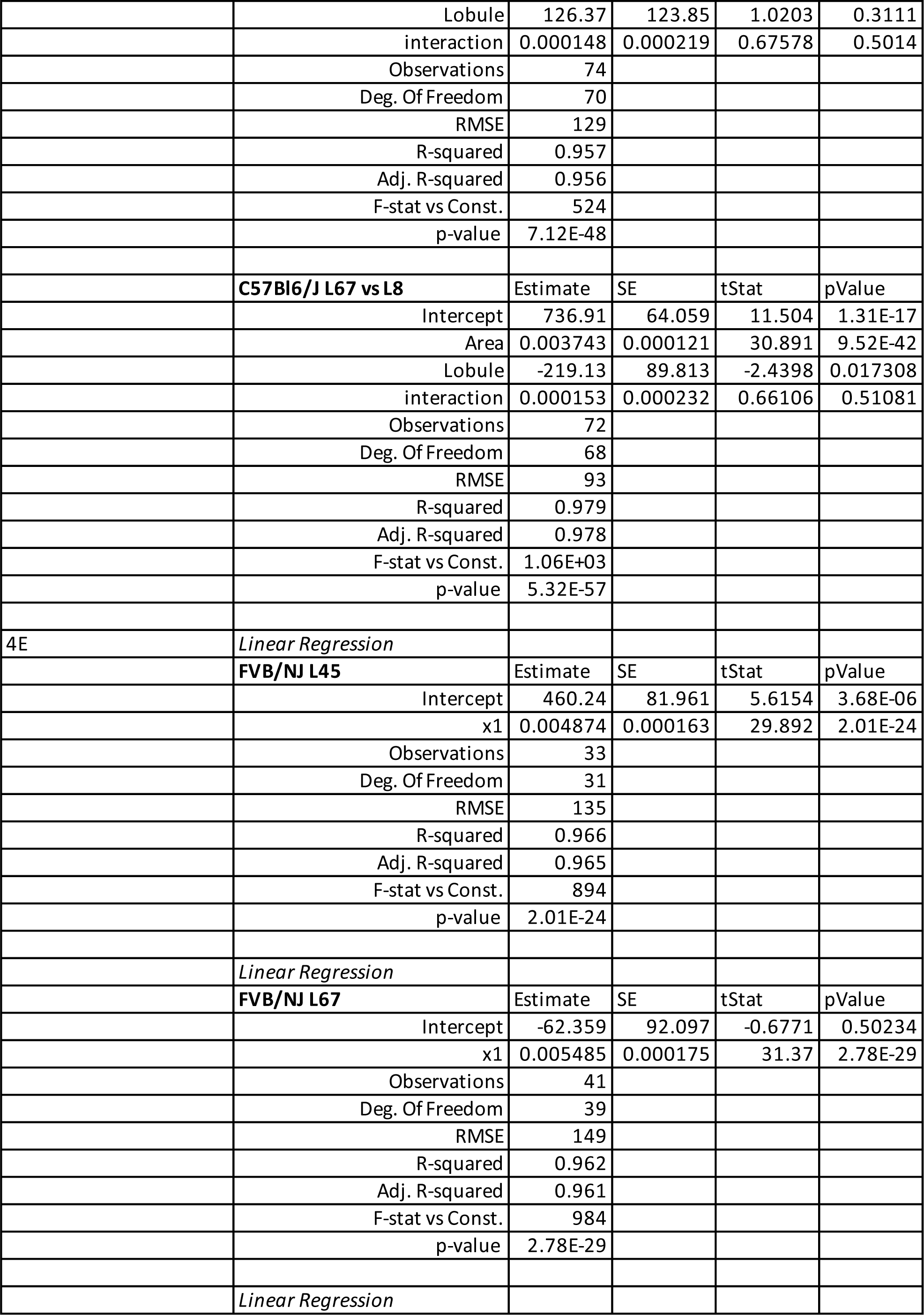
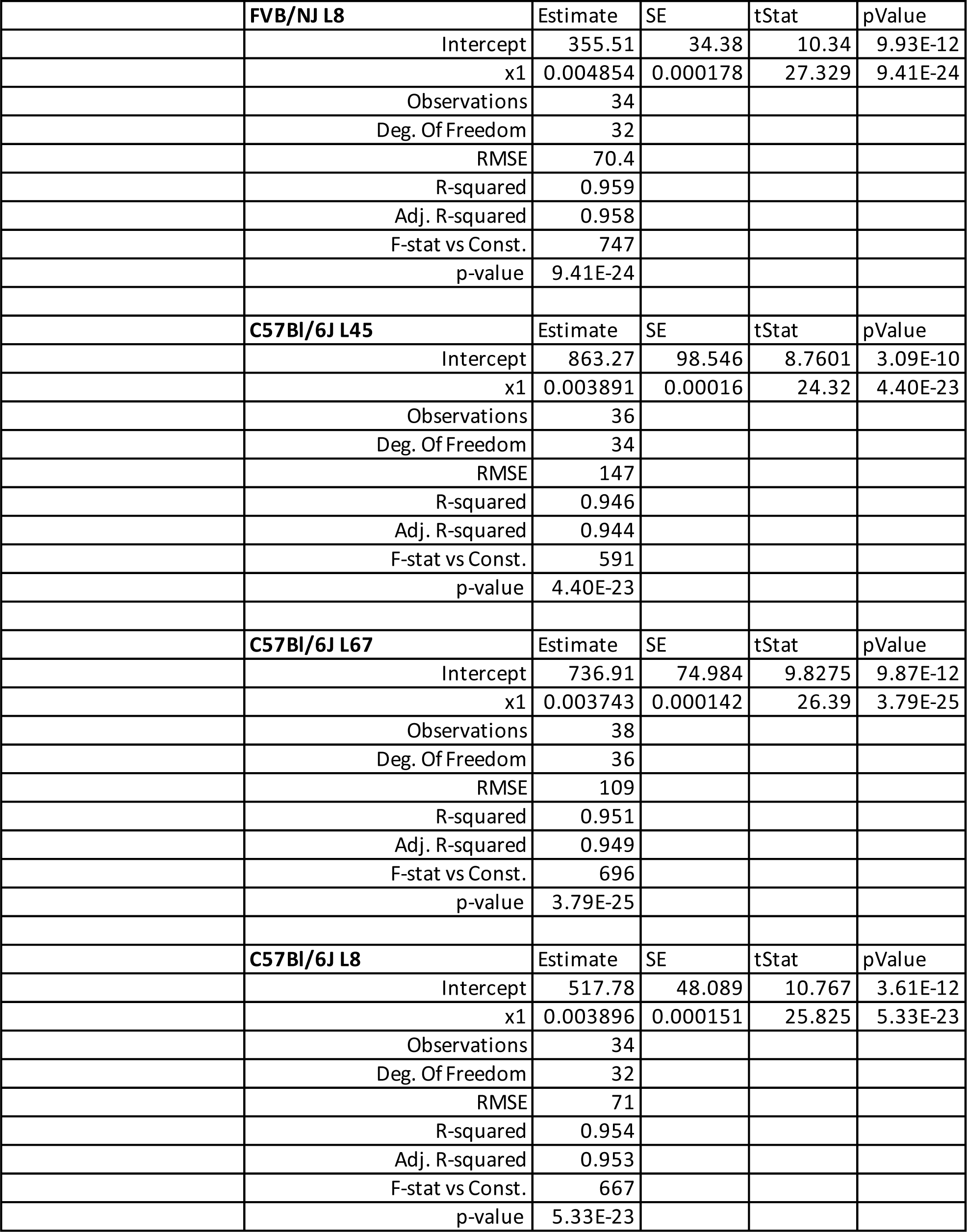
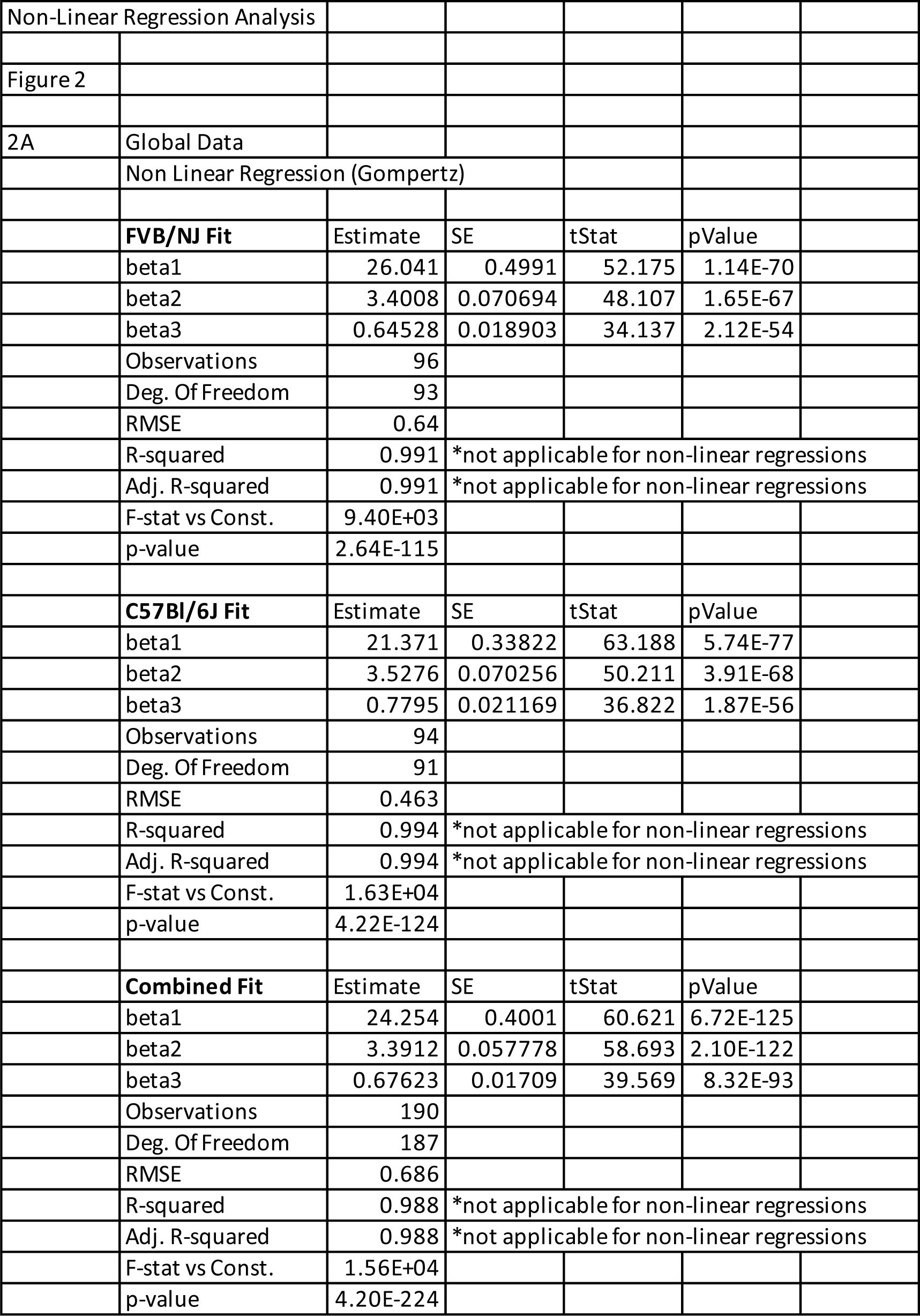

|  |  |  |  |  |  |
| --- | --- | --- | --- | --- | --- |
| Linear Regression Analysis |  |  |  |  |  |
| Main Figures |  |  |  |  |  |
| **Figure 3** |  |  |  |  |  |
| 3A | *Multiple Linear Regression* |  |  |  |  |
|  | **C57Bl6/J vs FVB/NJ** | Estimate | SE | tStat | pValue |
|  | Intercept | 736.91 | 90.476 | 8.1447 | 6.33E-12 |
|  | Area | 0.003743 | 0.000171 | 21.871 | 2.79E-34 |
|  | Strain | -799.26 | 121.51 | -6.5775 | 5.69E-11 |
|  | Interaction | 0.001742 | 0.00023 | 7.5644 | 8.02E-11 |
|  | Observations | 79 |  |  |  |
|  | Deg. Of Freedom | 75 |  |  |  |
|  | RMSE | 131 |  |  |  |
|  | R-squared | 0.959 |  |  |  |
|  | Adj. R-squared | 0.957 |  |  |  |
|  | F-stat vs Const. | 584 |  |  |  |
|  | p-value | 6.71E-52 |  |  |  |
| **Figure 6** |  |  |  |  |  |
| 6H | *Linear Regression* |  |  |  |  |
|  | **CDA vs Expansion** | Estimate | SE | tStat | pValue |
|  | Intercept | 226.22 | 45.036 | 5.0231 | 0.00737 |
|  | x1 | 374.56 | 60.287 | 6.213 | 0.003415 |
|  | Observations | 6 |  |  |  |
|  | Deg. Of Freedom | 4 |  |  |  |
|  | RMSE | 45.8 |  |  |  |
|  | R-squared | 0.906 |  |  |  |
|  | Adj. R-squared | 0.883 |  |  |  |
|  | F-stat vs Const. | 38.6 |  |  |  |
|  | p-value | 0.00342 |  |  |  |
| 6I | *Linear Regression* |  |  |  |  |
|  | **CDA vs Thickness** | Estimate | SE | tStat | pValue |
|  | Intercept | 56.859 | 3.257 | 17.458 | 8.07E-09 |
|  | x1 | -14.13 | 4.7075 | -3.0016 | 0.013308 |
|  | Observations | 12 |  |  |  |
|  | Deg. Of Freedom | 10 |  |  |  |
|  | RMSE | 4.24 |  |  |  |
|  | R-squared | 0.474 |  |  |  |
|  | Adj. R-squared | 0.421 |  |  |  |
|  | F-stat vs Const. | 9.01 |  |  |  |
|  | p-value | 0.0133 |  |  |  |
| Supplemental Figures |  |  |  |  |  |
| **Supplemental Fig 3** |  |  |  |  |  |
| 3I |  |  |  |  |  |
|  | *Linear Regression* |  |  |  |  |
|  | **FVB/NJ** | Estimate | SE | tStat | pValue |
|  | Intercept | 0.099801 | 0.10005 | 0.99751 | 0.32536 |
|  | x1 | 4.0451 | 0.17307 | 23.373 | 6.08E-23 |
|  | Observations | 37 |  |  |  |
|  | Deg. Of Freedom | 35 |  |  |  |
|  | RMSE | 0.211 |  |  |  |
|  | R-squared | 0.94 |  |  |  |
|  | Adj. R-squared | 0.938 |  |  |  |
|  | F-stat vs Const. | 546 |  |  |  |
|  | p-value | 6.08E-23 |  |  |  |
|  | *Linear Regression* |  |  |  |  |
|  | **C57Bl/6J** | Estimate | SE | tStat | pValue |
|  | Intercept | 0.06403 | 0.10781 | 0.59393 | 0.55627 |
|  | x1 | 3.9523 | 0.16637 | 23.755 | 1.37E-23 |
|  | Observations | 38 |  |  |  |
|  | Deg. Of Freedom | 36 |  |  |  |
|  | RMSE | 0.225 |  |  |  |
|  | R-squared | 0.94 |  |  |  |
|  | Adj. R-squared | 0.938 |  |  |  |
|  | F-stat vs Const. | 564 |  |  |  |
|  | p-value | 1.37E-23 |  |  |  |
|  | *Multiple Linear Regression* |  |  |  |  |
|  | **C57Bl6/J vs FVB/NJ** | Estimate | SE | tStat | pValue |
|  | Intercept | 0.06403 | 0.10441 | 0.61328 | 0.54165 |
|  | Area | 3.9523 | 0.16113 | 24.529 | 2.14E-36 |
|  | Strain | 0.035771 | 0.14711 | 0.24315 | 0.80859 |
|  | interaction | 0.092779 | 0.24104 | 0.38491 | 0.70146 |
|  | Observations | 75 |  |  |  |
|  | Deg. Of Freedom | 71 |  |  |  |
|  | RMSE | 0.218 |  |  |  |
|  | R-squared | 0.941 |  |  |  |
|  | Adj. R-squared | 0.938 |  |  |  |
|  | F-stat vs Const. | 374 |  |  |  |
|  | p-value | 1.99E-43 |  |  |  |
| **Supplemental Fig 4** |  |  |  |  |  |
| 4A | *Multiple Linear Regression* |  |  |  |  |
|  | **C57Bl6/J vs FVB/NJ L45** | Estimate | SE | tStat | pValue |
|  | Intercept | 863.27 | 94.726 | 9.1133 | 3.13E-13 |
|  | Area | 0.003891 | 0.000154 | 25.301 | 2.32E-35 |
|  | Strain | -403.03 | 127.89 | -3.1513 | 0.002458 |
|  | interaction | 0.000983 | 0.00023 | 4.275 | 6.39E-05 |
|  | Observations | 69 |  |  |  |
|  | Deg. Of Freedom | 65 |  |  |  |
|  | RMSE | 142 |  |  |  |
|  | R-squared | 0.96 |  |  |  |
|  | Adj. R-squared | 0.959 |  |  |  |
|  | F-stat vs Const. | 525 |  |  |  |
|  | p-value | 1.74E-45 |  |  |  |
| 4B | *Multiple Linear Regression* |  |  |  |  |
|  | **C57Bl6/J vs FVB/NJ L8** | Estimate | SE | tStat | pValue |
|  | Intercept | 517.78 | 47.879 | 10.814 | 4.41E-16 |
|  | Area | 0.003896 | 0.00015 | 25.939 | 1.14E-35 |
|  | Strain | -162.27 | 59.033 | -2.7489 | 0.007763 |
|  | interaction | 0.000958 | 0.000233 | 4.1063 | 0.000116 |
|  | Observations | 68 |  |  |  |
|  | Deg. Of Freedom | 64 |  |  |  |
|  | RMSE | 70.7 |  |  |  |
|  | R-squared | 0.972 |  |  |  |
|  | Adj. R-squared | 0.971 |  |  |  |
|  | F-stat vs Const. | 736 |  |  |  |
|  | p-value | 1.55E-49 |  |  |  |
| 4C | *Multiple Linear Regression* |  |  |  |  |
|  | **FVB/NJ L67 vs L45** | Estimate | SE | tStat | pValue |
|  | Intercept | -62.359 | 88.378 | -0.7056 | 0.48278 |
|  | Area | 0.005485 | 0.000168 | 32.69 | 3.95E-44 |
|  | Lobule | 522.6 | 123.86 | 4.2191 | 7.23E-05 |
|  | interaction | -0.00061 | 0.000241 | -2.5364 | 0.013432 |
|  | Observations | 74 |  |  |  |
|  | Deg. Of Freedom | 70 |  |  |  |
|  | RMSE | 143 |  |  |  |
|  | R-squared | 0.964 |  |  |  |
|  | Adj. R-squared | 0.962 |  |  |  |
|  | F-stat vs Const. | 623 |  |  |  |
|  | p-value | 2.10E-50 |  |  |  |
|  | **FVB/NJ L67 vs L8** | Estimate | SE | tStat | pValue |
|  | Intercept | -62.359 | 74.239 | -0.83998 | 0.40374 |
|  | Area | 0.005485 | 0.000141 | 38.916 | 1.26E-49 |
|  | Lobule | 417.86 | 94.636 | 4.4155 | 3.53E-05 |
|  | interaction | -0.00063 | 0.000334 | -1.8862 | 0.063362 |
|  | Observations | 75 |  |  |  |
|  | Deg. Of Freedom | 71 |  |  |  |
|  | RMSE | 120 |  |  |  |
|  | R-squared | 0.985 |  |  |  |
|  | Adj. R-squared | 0.984 |  |  |  |
|  | F-stat vs Const. | 1.55E+03 |  |  |  |
|  | p-value | 1.26E-64 |  |  |  |
| 4D | *Multiple Linear Regression* |  |  |  |  |
|  | **C57Bl6/J L67 vs L45** | Estimate | SE | tStat | pValue |
|  | Intercept | 736.91 | 88.878 | 8.2912 | 5.31E-12 |
|  | Area | 0.003743 | 0.000168 | 22.265 | 1.75E-33 |
|  | Lobule | 126.37 | 123.85 | 1.0203 | 0.3111 |
|  | interaction | 0.000148 | 0.000219 | 0.67578 | 0.5014 |
|  | Observations | 74 |  |  |  |
|  | Deg. Of Freedom | 70 |  |  |  |
|  | RMSE | 129 |  |  |  |
|  | R-squared | 0.957 |  |  |  |
|  | Adj. R-squared | 0.956 |  |  |  |
|  | F-stat vs Const. | 524 |  |  |  |
|  | p-value | 7.12E-48 |  |  |  |
|  | **C57Bl6/J L67 vs L8** | Estimate | SE | tStat | pValue |
|  | Intercept | 736.91 | 64.059 | 11.504 | 1.31E-17 |
|  | Area | 0.003743 | 0.000121 | 30.891 | 9.52E-42 |
|  | Lobule | -219.13 | 89.813 | -2.4398 | 0.017308 |
|  | interaction | 0.000153 | 0.000232 | 0.66106 | 0.51081 |
|  | Observations | 72 |  |  |  |
|  | Deg. Of Freedom | 68 |  |  |  |
|  | RMSE | 93 |  |  |  |
|  | R-squared | 0.979 |  |  |  |
|  | Adj. R-squared | 0.978 |  |  |  |
|  | F-stat vs Const. | 1.06E+03 |  |  |  |
|  | p-value | 5.32E-57 |  |  |  |
| 4E | *Linear Regression* |  |  |  |  |
|  | **FVB/NJ L45** | Estimate | SE | tStat | pValue |
|  | Intercept | 460.24 | 81.961 | 5.6154 | 3.68E-06 |
|  | x1 | 0.004874 | 0.000163 | 29.892 | 2.01E-24 |
|  | Observations | 33 |  |  |  |
|  | Deg. Of Freedom | 31 |  |  |  |
|  | RMSE | 135 |  |  |  |
|  | R-squared | 0.966 |  |  |  |
|  | Adj. R-squared | 0.965 |  |  |  |
|  | F-stat vs Const. | 894 |  |  |  |
|  | p-value | 2.01E-24 |  |  |  |
|  | *Linear Regression* |  |  |  |  |
|  | **FVB/NJ L67** | Estimate | SE | tStat | pValue |
|  | Intercept | -62.359 | 92.097 | -0.6771 | 0.50234 |
|  | x1 | 0.005485 | 0.000175 | 31.37 | 2.78E-29 |
|  | Observations | 41 |  |  |  |
|  | Deg. Of Freedom | 39 |  |  |  |
|  | RMSE | 149 |  |  |  |
|  | R-squared | 0.962 |  |  |  |
|  | Adj. R-squared | 0.961 |  |  |  |
|  | F-stat vs Const. | 984 |  |  |  |
|  | p-value | 2.78E-29 |  |  |  |
|  | *Linear Regression* |  |  |  |  |

|  | **FVB/NJ L8** | Estimate | SE | tStat | pValue |
| --- | --- | --- | --- | --- | --- |
|  | Intercept | 355.51 | 34.38 | 10.34 | 9.93E-12 |
|  | x1 | 0.004854 | 0.000178 | 27.329 | 9.41E-24 |
|  | Observations | 34 |  |  |  |
|  | Deg. Of Freedom | 32 |  |  |  |
|  | RMSE | 70.4 |  |  |  |
|  | R-squared | 0.959 |  |  |  |
|  | Adj. R-squared | 0.958 |  |  |  |
|  | F-stat vs Const. | 747 |  |  |  |
|  | p-value | 9.41E-24 |  |  |  |
|  | **C57Bl/6J L45** | Estimate | SE | tStat | pValue |
|  | Intercept | 863.27 | 98.546 | 8.7601 | 3.09E-10 |
|  | x1 | 0.003891 | 0.00016 | 24.32 | 4.40E-23 |
|  | Observations | 36 |  |  |  |
|  | Deg. Of Freedom | 34 |  |  |  |
|  | RMSE | 147 |  |  |  |
|  | R-squared | 0.946 |  |  |  |
|  | Adj. R-squared | 0.944 |  |  |  |
|  | F-stat vs Const. | 591 |  |  |  |
|  | p-value | 4.40E-23 |  |  |  |
|  | **C57Bl/6J L67** | Estimate | SE | tStat | pValue |
|  | Intercept | 736.91 | 74.984 | 9.8275 | 9.87E-12 |
|  | x1 | 0.003743 | 0.000142 | 26.39 | 3.79E-25 |
|  | Observations | 38 |  |  |  |
|  | Deg. Of Freedom | 36 |  |  |  |
|  | RMSE | 109 |  |  |  |
|  | R-squared | 0.951 |  |  |  |
|  | Adj. R-squared | 0.949 |  |  |  |
|  | F-stat vs Const. | 696 |  |  |  |
|  | p-value | 3.79E-25 |  |  |  |
|  | **C57Bl/6J L8** | Estimate | SE | tStat | pValue |
|  | Intercept | 517.78 | 48.089 | 10.767 | 3.61E-12 |
|  | x1 | 0.003896 | 0.000151 | 25.825 | 5.33E-23 |
|  | Observations | 34 |  |  |  |
|  | Deg. Of Freedom | 32 |  |  |  |
|  | RMSE | 71 |  |  |  |
|  | R-squared | 0.954 |  |  |  |
|  | Adj. R-squared | 0.953 |  |  |  |
|  | F-stat vs Const. | 667 |  |  |  |
|  | p-value | 5.33E-23 |  |  |  |
| Non-Linear Regression Analysis |  |  |  |  |  |
| Figure 2 |  |  |  |  |  |
| 2A | Global Data |  |  |  |  |
|  | Non Linear Regression (Gompertz) |  |  |  |  |
|  | **FVB/NJ Fit** | Estimate | SE | tStat | pValue |
|  | beta1 | 26.041 | 0.4991 | 52.175 | 1.14E-70 |
|  | beta2 | 3.4008 | 0.070694 | 48.107 | 1.65E-67 |
|  | beta3 | 0.64528 | 0.018903 | 34.137 | 2.12E-54 |
|  | Observations | 96 |  |  |  |
|  | Deg. Of Freedom | 93 |  |  |  |
|  | RMSE | 0.64 |  |  |  |
|  | R-squared | 0.991 | *not applicable for non-linear regressions |  |  |
|  | Adj. R-squared | 0.991 | *not applicable for non-linear regressions |  |  |
|  | F-stat vs Const. | 9.40E+03 |  |  |  |
|  | p-value | 2.64E-115 |  |  |  |
|  | **C57Bl/6J Fit** | Estimate | SE | tStat | pValue |
|  | beta1 | 21.371 | 0.33822 | 63.188 | 5.74E-77 |
|  | beta2 | 3.5276 | 0.070256 | 50.211 | 3.91E-68 |
|  | beta3 | 0.7795 | 0.021169 | 36.822 | 1.87E-56 |
|  | Observations | 94 |  |  |  |
|  | Deg. Of Freedom | 91 |  |  |  |
|  | RMSE | 0.463 |  |  |  |
|  | R-squared | 0.994 | *not applicable for non-linear regressions |  |  |
|  | Adj. R-squared | 0.994 | *not applicable for non-linear regressions |  |  |
|  | F-stat vs Const. | 1.63E+04 |  |  |  |
|  | p-value | 4.22E-124 |  |  |  |
|  | **Combined Fit** | Estimate | SE | tStat | pValue |
|  | beta1 | 24.254 | 0.4001 | 60.621 | 6.72E-125 |
|  | beta2 | 3.3912 | 0.057778 | 58.693 | 2.10E-122 |
|  | beta3 | 0.67623 | 0.01709 | 39.569 | 8.32E-93 |
|  | Observations | 190 |  |  |  |
|  | Deg. Of Freedom | 187 |  |  |  |
|  | RMSE | 0.686 |  |  |  |
|  | R-squared | 0.988 | *not applicable for non-linear regressions |  |  |
|  | Adj. R-squared | 0.988 | *not applicable for non-linear regressions |  |  |
|  | F-stat vs Const. | 1.56E+04 |  |  |  |
|  | p-value | 4.20E-224 |  |  |  |

## Notes

### Competing Interest Statement

The authors have declared no competing interest.

